# Decisive Roles of Sequence Distributions in the Generalizability of *de novo* Deep Learning Models for RNA Secondary Structure Prediction

**DOI:** 10.1101/2022.06.29.498185

**Authors:** Xiangyun Qiu

## Abstract

Taking sequences as the only inputs, the class of *de novo* deep learning (DL) models for RNA secondary structure prediction has achieved far superior performances than traditional algorithms. However, key questions remain over the statistical underpinning of such models that make no use of physical laws or co-evolutionary information. We present a quantitative study of the capacity and generalizability of a series of *de novo* DL models, with a minimal two-module architecture and no post-processing, under varied distributions of the seen and unseen sequences. Our DL models outperform existing methods on commonly used benchmark datasets and demonstrate excellent learning capacities under all sequence distributions. These DL models generalize well over non-identical unseen sequences, but the generalizability degrades rapidly as the sequence distributions of the seen and unseen datasets become dissimilar. Examinations of RNA family-specific behaviors manifest not only disparate family-dependent performances but substantial generalization gaps within the same family. We further determine how model generalization decreases with the decrease of sequence similarity via pairwise sequence alignment, providing quantitative insights into the limitations of statistical learning. Model generalizability thus poses a major hurdle for practical uses of *de novo* DL models and several tenable avenues for future advances are discussed.

## INTRODUCTION

As a linear chain of nucleotides capable of base pairing, an RNA molecule readily forms various secondary structure motifs such as stems and loops, regardless of the existence of stable tertiary structures (1–3). Particularly for the diverse families of non-coding RNAs (ncRNA) (4), their secondary structures are more conserved than sequences and provide important cues for their biological functions (5,6). And even messenger RNAs (mRNA) possess key secondary structure motifs for translation initiation and regulation (7–9). As such, there have been major interests in determining and understanding RNA secondary structures, via both experiment and computation (10–12). In recent years, with the emergence of sizeable RNA structure databases and the accessibility of powerful machine learning algorithms, data-centric deep-learning-based models, the subject of this study, have been successfully developed for RNA secondary structure prediction (13).

Predicting RNA secondary structures comes down to the identification of all native base pairs. As any two nucleotides (i.e., AUCG) can theoretically pair up (14), the native set of base pairs is regarded as the optimal among all possible sets. Such an optimal set is best inferred from the covariance patterns of homologous sequences which can be costly or impossible to obtain (10,15). In lieu of homologous sequences, traditional *de novo* computational approaches generally represent the secondary structure as a graph with nucleotides as nodes and base pairs as edges. A score is then computed according to pre-defined structural elements of the graph, using parameters derived from physical measurements of thermodynamic energies (16), data mining of known RNA secondary structures, or a combination of both (17). The onus for the predictive task is to optimize the score by searching the entire structure space, which however grows exponentially with the RNA sequence length. In order to reduce the computational complexity, various rules or constraints of RNA secondary structures have been introduced, such as non-nested base pairing (i.e., no pseudoknots), canonical base pairs only (i.e., AU, GC, and GU), and no sharp turns (i.e., loop length > 3). Varieties of efficient algorithms, especially based on dynamic programming and related techniques (18), have also been introduced along with improved scoring parameters (16,17). However, traditional algorithms have struggled to make significant gains in performance in the recent decade (13).

A major recent advance in prediction performance comes from the application of deep learning (DL) to the RNA secondary structure problem. Instead of the graph search in traditional methods, DL models represent the secondary structure as a 2D pairing probably matrix (PPM) and directly predict every PPM*_ij_* (*i* and *j* are indices to the nucleotides) in a parallel and quasi-independent manner. In order to directly map an input sequence to its predicted PPM, DL models often employ many abstraction layers enlisting a large number of parameters that must be learned via training on known RNA secondary structures. As existing RNA secondary structure datasets are curated largely via comparative sequence analysis (19,20), this study focuses on the class of single-sequence-based DL models (i.e., without the use of co-evolutionary sequences), also referred to as *de novo* DL models herein, noting that homologous-sequence modeling with tools such as TurboFold (21) and R-scape (22) is still expected to the most accurate. Several recent publications demonstrated that such *de novo* DL models can be successfully developed with up to several millions of parameters, such as 2dRNA (23), ATTfold (24), DMfold (25), E2Efold (26), MXfold2 (27), SPOT-RNA (28), and Ufold (29), among others (30–33). These DL models substantially outperform traditional algorithms, with even close-to-perfect predictions for certain benchmarking datasets (e.g., Ufold on the ArchiveII dataset (34)). As a result, single-sequence-based DL models have emerged as a promising and powerful solution to the RNA secondary structure problem.

Despite the recent successes, important questions exist for practical uses of such *de novo* DL models. One is that existing *de novo* DL models have yet to reach satisfactory performances on all known datasets. For example, the best F1 score for the largest dataset, bpRNA (19), remains low, ~0.65, by all DL and traditional models. And this best score is attained by Ufold after incorporation of prior knowledge of RNA structure, specifically with an iterative post-processing module to enforce structural constraints and sparsity. This calls into the question of the learning capacity of *de novo* DL models given the considerable size of 8.6M parameters in the largest model (Ufold). Another related, arguably more critical, issue concerns the generalizability of the *de novo* DL models, i.e., how DL models perform over an unseen/test set of RNA sequences compared with the seen/training set. Substantial performance drops would indicate poor generalizability to which DL models are highly susceptible. To mitigate such tendency, various strategies have been explored, e.g., E2Efold and Ufold enforce known structural constraints via iterative refinement, SPOT-RNA uses an ensemble of five models, and MXfold2 integrates DL-predicted folding scores with thermodynamic regulation. Nonetheless, the generalizability of current DL models strongly depends on the sequence similarities between the seen and unseen datasets (35) and model comparisons are only carried out when the same pair of seen and unseen datasets are used. While this issue is likely inherent to all DL models because of their data-centric statistical learning, such non-data agnostic generalizability presents a major hurdle for practical uses of *de novo* DL models for RNA secondary structure prediction.

To address these questions, we have investigated the capacity and generalizability of *de novo* DL models with different model sizes and under varied sequence distributions of the seen/unseen datasets. We chose a minimal two-module network architecture without any post processing (except for discretization with the threshold of 0.5) so as to probe the intrinsic behaviors of DL models. We found that a rather small DL model of 16K parameters can achieve decent performances over a medium-sized dataset and that medium-sized models with less than 1M parameters can attain excellent performances and surpass existing DL or traditional models. However, model generalization deteriorates as the sequence similarity between the seen and unseen datasets decreases. In order to gain quantitative insights into model generalizability, we determined how model generalization depends on the similarity between the seen and unseen sequences via pairwise sequence alignment. Our observations indicate that *de novo* DL models appear to be largely statistical learners of the correlation between RNA sequences and their secondary structures and we last discuss various pathways to improve model generalizability and develop data-agonistic *de novo* DL models.

## MATERIALS AND METHODS

Two main ingredients of this study are the types of DL network architecture and the sequence distributions of the seen and unseen datasets. We reason that, given an RNA sequence, each nucleotide first explores all its molecular surroundings and then engages in the dynamic process of paring and unpairing before arriving at the most stable configuration. A two-module network architecture was thus chosen to operate at the sequence and pair levels, respectively. To evaluate model capacity and generalizability, we chose three similarity levels between the seen and unseen datasets as follows. The first level only requires no identical sequences between the seen and unseen sets, i.e., the cross-sequence level. The second level further stipulates that all pairs of sequences in both datasets are below 80% in sequence identity (filtered with the software CD-HIT (36)), referred to as the cross-cluster level. The third level is the most stringent by having the seen and unseen sets from different RNA families, e.g., one from tRNA and the other from 5S rRNA, named as the cross-family level. Brief descriptions of the datasets and the model development are given below and more details are provided in the Suppl. Materials.

### Datasets

With the three similarity levels of sequence distributions in mind, we choose to primarily work with two recent datasets, RNA Stralign curated in 2017 (21) and ArchiveII curated in 2016 (34), both of which are medium-sized, comprehensive databases developed for benchmarking RNA secondary structure predictions. Compared with the only other larger comprehensive collection (bpRNA (19), also used in this study), RNA family types are conveniently provided by Stralign and ArchiveII, facilitating intra- and inter-family examinations. Both datasets have also been used by several DL models (e.g., E2Efold, MXfold2, and Ufold) with pre-trained model parameters available. For training efficiency and consistency with other DL models, RNA sequences longer than 600 nucleotides are excluded. For the cross-sequence level, only duplicated sequences are deleted from each dataset, yielding Stralign ND-L600 and ArchiveII ND-L600 with 20,118 and 3,395 sequences, respectively. CD-HIT-EST is then used to remove sequences with over 80% sequence identity, yielding the Stralign NR80-L600 and ArchiveII NR80-L600 datasets for the cross-cluster study. The ArchiveII NR80-L600 sequences with over 80% identity to Stralign NR80-L600 are further removed to give the ArchiveII-Stralign NR80-L600 dataset. Fig. 1A shows the population distributions of the eight RNA families of the Stralign dataset at different sequence similarity levels, evidencing vastly different redundancies for different families. For the cross-family study, the Stralign and ArchiveII datasets are combined to create the Strive dataset, which is processed in the same way to obtain its ND-L600 and NR80-L600 subsets. More information on these and other datasets (e.g., bpRNA) is given in the Suppl. Materials.

**Figure 1.**
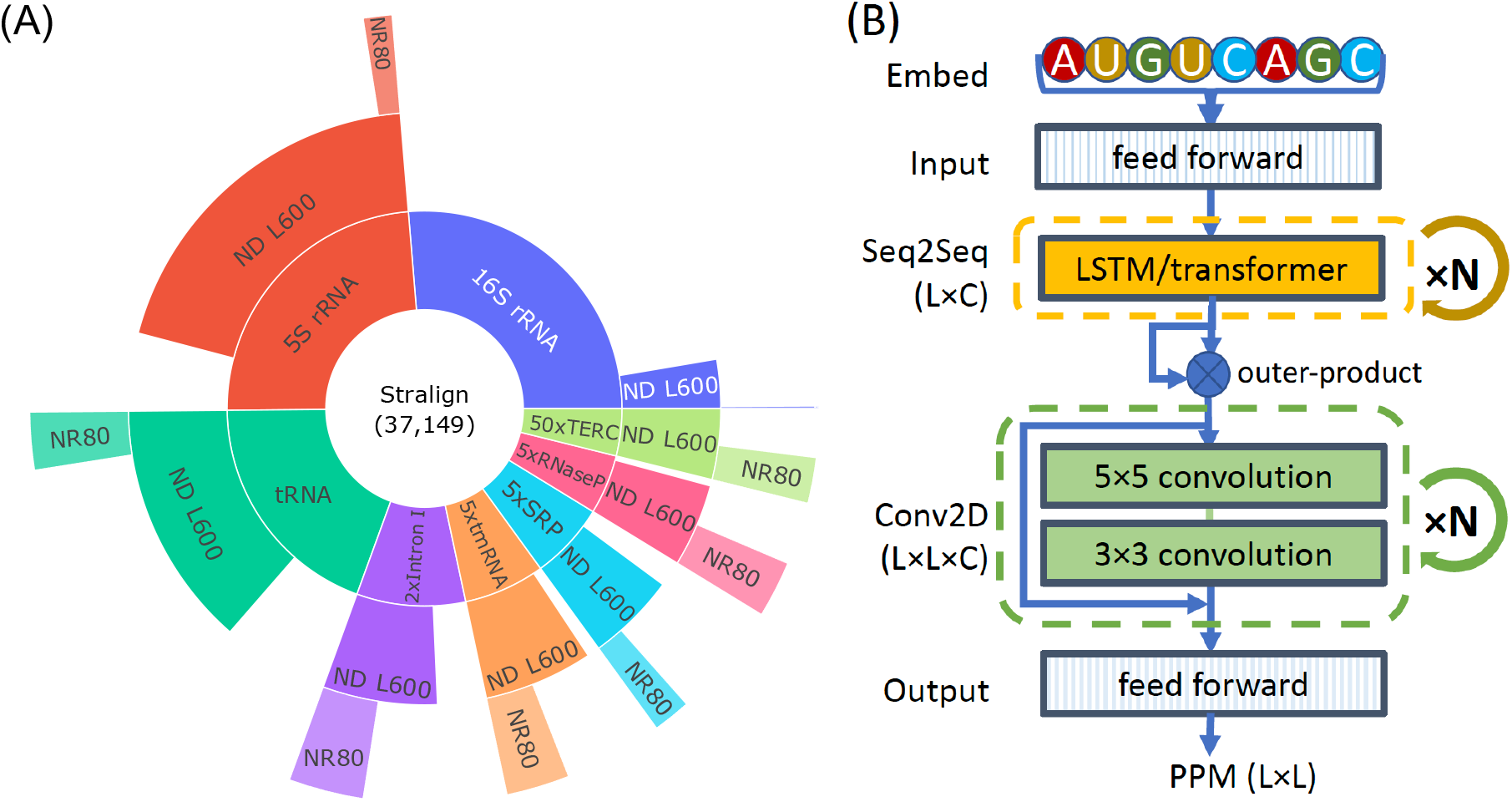
Illustrations of the Stralign dataset and the SeqFold2D architecture. (A) The population distributions of eight RNA families in the Stralign dataset at different sequence redundancy levels. The abbreviations of RNA families are, rRNA: ribosomal RNA, tRNA: transfer RNA, Intron I: group I intron, tmRNA: transfer messenger RNA, SRP: signal recognition particle, and TERC: telomerase RNA component. The innermost ring shows the population distributions of the RNA families in the original Stralign dataset with a total of 37,149 sequences, noting that the five less represented families (counter-clockwise from Intron I to TERC) are scaled up for visibility and the multiplier N is shown as “N×” in the label (see Suppl. Fig. S1 for the unscaled version). The middle ring shows the population of each RNA family type at the cross-sequence level after removing duplicated sequences and sequences longer than 600 nucleotides, labelled as “ND L600”. The outermost ring shows RNA family populations at the cross-cluster level after removing redundant sequences with 80% sequence identity or higher, labelled as “NR80”. Note that the number of sequences for the 16S rRNA NR80 set is only 50 and hardly visible. (B) The two-module network diagram of the SeqFold2D models. An input RNA sequence of length L is first one-hot encoded and stacked as an L×12 tensor for k-mer (k=3) representation, followed by feed-forward layers to yield an L×C tensor. The first main module consists of N stacked blocks of either bidirectional Long-Short-Term-Memory (LSTM) or transformer encoders. The resultant L×C tensor is then transformed into the L×L×C pair representation via outer-product, before being fed to the second main module of N stacked blocks of residual 2D convolutional layers. The output block is made up of three feed-forward layers and predicts the pairing probability matrix (PPM) of dimension L×L.

### Network architecture

As shown in Fig. 1B, our DL neural network, named SeqFold2D, comprises two main learning modules flanked by the input and output blocks. As the only input to the SeqFold2D network, each RNA sequence of length L is first one-hot-encoded as an L×4 tensor and then stacked into its k-mer (k=3) representation of L×12. The input block mixes the twelve channels into an L×C tensor with two feed-forward layers. The channel size C is then kept constant for the rest of the network unless noted otherwise. The first main learning module is made up of N repeated blocks of bidirectional Long-Short-Term-Memory (LSTM) or selfattention-based transformer encoder layers to learn richer representations from the whole sequence, which are then transformed into 2D pair representations (L×L×C) via outer-product. The second learning module consists of the same number (N) of repeated blocks of residual 2D convolutional layers to learn pair interactions locally. The output block consists of three feedforward layers with the channel size C=2 for the final layer. Softmax is then used to yield the continuous PPM of shape L×L. Non-linear activations (LeakyReLU or Swish) are applied before every linear operation with weights and biases, followed by layer normalizations and dropouts (0.2-0.42). The size of a specific SeqFold2D network is thus determined by two design variables, N (the number of blocks) and C (the channel size). For example, N=1 and C=16 gives ~16K trainable parameters and N=4 and C=64 gives ~960K parameters.

### Performance metrics and benchmarking

We use the F1 score as the main metric for evaluating model performances (37). It is the harmonic mean of Precision and Recall and defined as 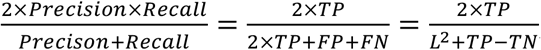, where Precision is TP/(TP+FP), Recall TP/(TP+FN), TP the number of true positives, FP false positives, FN false negatives, and TN true negatives. In order to examine the exclusive capabilities of DL models, no post-processing is applied to the continuous PPM except for discretization with the threshold of 0.5, i.e., no grid search, model ensemble averaging, thermodynamic regulation, or enforcement of RNA structure constraints. In addition to several *de novo* DL models, we further benchmark the SeqFold2D models against the following traditional algorithms: Mfold/UNAfold (38), RNAfold (39), RNAstructure(40), LinearFold (41), SimFold (42), CONTRAfold (43), ContextFold(44), and Centroidfold (45). Note that we categorize machine-learning-based hybrid models as traditional algorithms for wording simplicity.

### Model development and evaluation of overfitting and generalizability

All SeqFold2D models were implemented with the Paddle framework (https://github.com/PaddlePaddle/Paddle) and trained with the AdamW optimizer in two stages. The first stage uses the binary cross-entropy (CE) loss between the predicted PPM and the ground truth with equal weights for positive and negative labels. Note that E2Efold and Ufold weighted the positive labels with a constant of 300 to address the issue of imbalanced negative/positive distributions, which we found to be helpful in the beginning but lead to higher ratios of false positives in the end. Once the CE loss plateaus, the second stage is invoked with the soft F1-score loss used by E2Efold and others.

The two-stage training was found to give the best F1 score compared with the use of either loss function alone. For hyperparameter tuning, we carried out limited manual searches for the learning rate, batch size, and dropout rate. No significant differences were observed between the uses of LSTM or transformer layers for the first module, though the LSTM layers led to more consistent training outcomes.

We refer to the set of sequences used to develop a specific SeqFold2D model as the seen set that consists of two subsets, training and validation. The training set is used to train model weights and biases and the validation set is used to optimize model hyperparameters such as the number of epochs, weight decay, and learning rate. We refer to the test set for benchmarking models also as the unseen set. These different sets of sequences are obtained via random splits stratified by the family type from their corresponding parent dataset. Model performances on the training (TR), validation (VL), and test (TS) sets can all be different, usually with the best for TR and the worst for TS.

In order to take a closer look into model behaviors, we further distinguish two different performance gaps. The first is between the TR and VL sets that are always taken from the same parent dataset in this study. We reason that the TR-VL variance indicates whether a DL model is learning both TR and VL sets (i.e., no overfitting) or additional spurious patterns of the TR set only (i.e., overfitting). The TR-VL variance is thus used as the indicator for model overfitting. The second is between the TR and TS sets that may have similar or different sequence distributions. Hence the TR-TS variance reflects model generalizability at the specific level of sequence similarity and is also referred to as the generalization gap. Importantly, every SeqFold2D model herein was trained to optimize the F1 score on its corresponding VL set, and the model parameters giving the highest VL F1 score were saved as the final model to be benchmarked on the unseen TS set. One alternative is to stop training as soon as the TR-VL variance becomes significant (e.g., 5%), which we chose not to follow given our purpose of investigating model capacity and generalizability.

## RESULTS

### Cross-sequence study: excellent capacity and generalizability of *de novo* DL models

We first use the larger Stralign ND-L600 dataset (Stral-ND in short) to train and test SeqFold2D models of various sizes at the cross-sequence level. The TR, VL, and TS sets are randomly split from Stral-ND at the ratios of 70%, 15%, and 15%, respectively. By varying the number of blocks (N) and the channel size (C), we gradually increase the number of model parameters from ~16K (N=1, C=16) to ~960K (N=4, C=64). Fig. 2A shows the F1 scores for the TR and TS sets from five SeqFold2D models and selected traditional models. Note that other DL models are not included here as they were trained with different dataset configurations to be examined next. As the model size increases, the performances of the SeqFold2D models increase steadily for both TR and TS and surpass all other algorithms by a large margin. Notably, the smallest SeqFold2D model with 16K parameters achieves an F1 score of ~0.8 and the largest model, SeqFold2D-960K, attains a nearly perfect performance (F1~0.985). Meanwhile, the TR-TS generalization gaps show to be negligible for the SeqFold2D models of sizes up to 420K parameters. A slight drop (~1.5%) in the F1 score can be spotted for the SeqFold2D-960K model (the top bars in Fig. 2A). As the F1-score distributions are non-Gaussian, we apply the Kolmogorov-Smirnov (KS) test and find the TR-TS variance to be indeed statistically significant for the 960K model (P-value around 3e-6) but insignificant for all other SeqFold2D models (P-values greater than 0.1).

**Figure 2.**
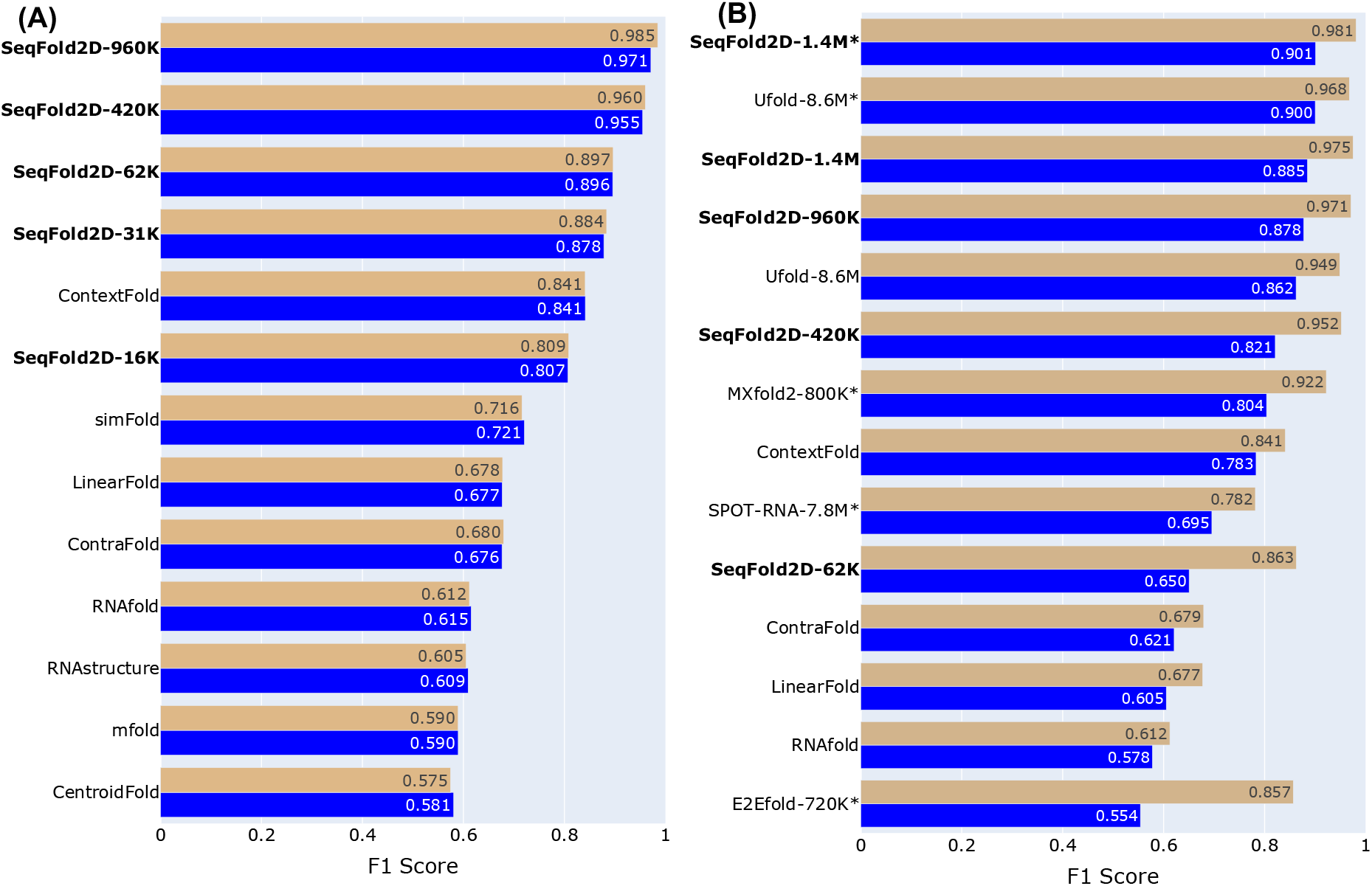
The F1 scores on the TR (tan) and TS (blue) sets by SeqFold2D models and other selected DL and traditional models in two different dataset setups. The number of parameters is shown after the name and the F1 value is given at the end of each bar. (A) Both TR and TS sets are taken from the Stralign ND-L600 dataset. (B) The TR set is taken from the Stralign ND-L600 dataset and the TS set is the entire ArchiveII ND-L600 dataset. The asterisk after the name indicates the use of post-processing.

Nonetheless, the TR-TS generalization gaps are rather small and we further verify that the TR-VL variances show similar behaviors (see Suppl. Fig. S8). Altogether, the SeqFold2D models demonstrate excellent expressive capacities for mapping RNA sequences to their secondary structures and generalize to high degrees of performances at the cross-sequence level (e.g., F1~0.97 on the TS set by SeqFold2D-960K).

In order to compare with other *de novo* DL models such as MXfold2 and Ufold, we next follow the same dataset setup of using the Stral-ND dataset as the TR and VL sets and the ArchiveII-ND-L600 (Archi-ND in short) as the TS set. Fig. 2B shows the F1 scores from several DL and traditional models, including five SeqFold2D models with 62K to 1.4M parameters. Note that the only available SPOT-RNA model was trained with the bpRNA dataset and its performance should not be directly compared. As previously reported for the Archi-ND dataset (26,27,29), all DL models, including the SeqFold2D models, outperform traditional algorithms substantially and attain F1 scores as high as ~0.9. Notably, both E2Efold and Ufold post-process predicted PPMs by excluding non-canonical base pairs and sharp turns and enforcing sparsity through iterative refinement. In part prompted by the superior performance of Ufold over all SeqFold2D models on the TS set, we examined the effectiveness of such post-processing by comparing the performances of Ufold with and without post-processing, shown as Ufold-8.6M* and Ufold-8.6M in Fig. 2B, respectively. Considerable gains in F1 scores were observed, ~2% for Stral-ND and ~4% for Archi-ND. We subsequently experimented with the same training and post-processing steps for the SeqFold2D-1.4M* model and realized similar performance gains, e.g., producing the best F1 scores for both Stral-ND (0.981) and Archi-ND (0.901) among all models. However, such post processing requires prior knowledge of the test set since the Archi-ND dataset indeed has canonical base pairs only and no sharp turns. We therefore did not use post-processing for all other SeqFold2D models. Overall, the SeqFold2D models show to compare favorably against other *de novo* DL models for both datasets, albeit with fewer parameters.

Another emphatic trend in Fig. 2B is that all DL models, including SeqFold2D, exhibit significant generalization gaps between Stral-ND and Archi-ND, with F1-score falloffs ranging from 8% to nearly 40%. This also holds true for MXfold2 with thermodynamic regularization and SPOT-RNA trained on different datasets. However, we observed very little TR-VL variances in training all SeqFold2D models (Suppl. Fig. S9) and also carried out five-fold cross-validation to rule out the likelihood of fortuitous data splitting. Therefore, the SeqFold2D models are not overfitting the TR set over the VL set but faithfully describing the entire Stral-ND distribution. Different sequence distributions between Stral-ND and Archi-ND are thus left as the most plausible cause for the observed generalization gap, which is corroborated by even larger performance drops for more dissimilar datasets such as bpRNA. On the whole, Stral-ND and Archi-ND have nearly identical RNA family types (the only exception is 23S rRNA in Archi-ND only, but at a mere 0.6%), but with very different population distributions of RNA families (Suppl. Fig. S5). This motivated us to examine the RNA family-specific performances of DL models, reasoning that Archi-ND may happen to have higher fractions of more difficult RNA families.

Fig. 3 shows the F1 scores per RNA family on Stral-ND and Archi-ND for the SeqFold2D-1.4M model. We indeed observe divergent family-specific model performances, for example, F1~0.998 for tRNA and F1~0.764 for telomerase RNA (TERC). The general trend is the better performances for the more populous families as expected from the statistical nature of DL model training. However, instead of similar family-wise performances between the seen and unseen, large drops within the same family are observed for most RNA families, with the largest being ~41% for 16S rRNA and a significant ~5% for tRNA (0.998 to 0.952). We further verified that another DL model (Ufold-8.6M*) manifests the same qualitative family-wise performance gaps (Suppl. Fig. S10). It is worth noting that two families (tmRNA and TERC) show almost no difference and our subsequent analyses indicate their sequences are highly redundant across Stral-ND and Archi-ND. In addition, the SeqFold2D-1.4M model gives nearly identical F1 scores between the Stral-ND TR and VL sets for all RNA families (Suppl. Fig. S11), ruling out the possibility that some RNA families are overfit and some are not. Altogether, these lead to a somewhat surprising observation that the *de novo* DL models are not guaranteed to generalize within the same RNA family type that is supposedly made up of closely related sequences.

**Figure 3.**
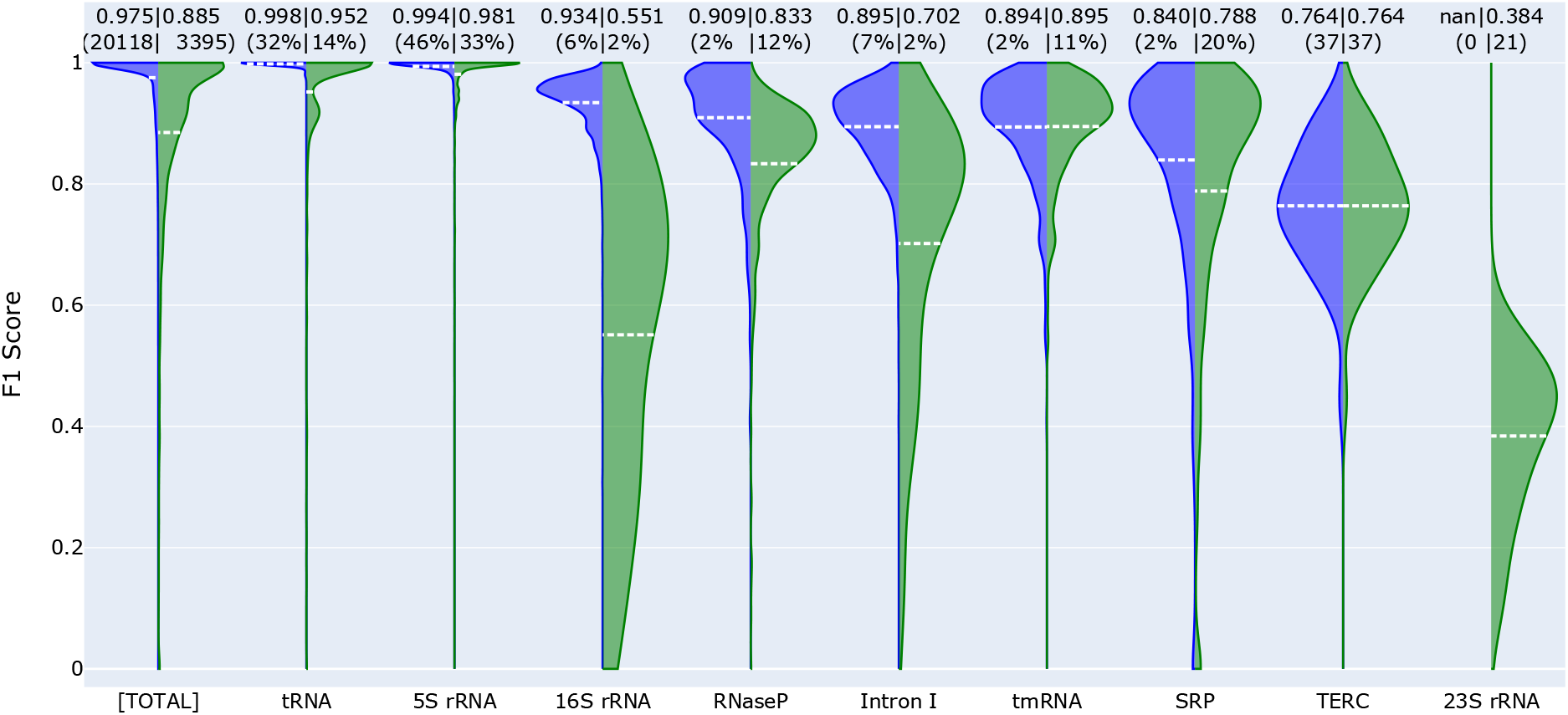
The F1 scores on the TR set taken from Stralign ND-L600 (left, blue) and the TS set of ArchiveII ND-L600 (right, green) by the SeqFold2D-1.4M model. The leftmost pair of violins show the F1 scores for the whole seen and unseen set and the following violin pairs show the F1 scores for each constituent RNA family. The averaged F1 scores are shown at the top and also as dashed lines (white) in the corresponding violins. The values in the parentheses above the violins are the sequence counts in actual numbers (for the whole set or families with less than 1% population shares) or in percentages (for families with more than 1% population shares). Note that 23S rRNA only exists in the ArchiveII ND-L600 dataset and is thus shown as nan for the Stralign ND-L600 dataset.

### Cross-cluster study: strong propensity towards overfitting and degraded generalizability

The sequence distributions within the same RNA family can be highly uneven when only identical sequences are removed (i.e., at the cross-sequence level). One way to mitigate such imbalance is to cluster similar sequences and remove all but one redundant sequence from each cluster. We applied this procedure using CD-HIT with the cluster similarity cutoff of 80%, the lowest allowed by CD-HIT, and obtained the non-redundant Stral-NR80 (3,122 RNAs) and Archi-Stral-NR80 (433 RNAs) datasets as aforementioned. Out of curiosity, we verified that all inter-family sequences are below 80% similarity level for all datasets. With Stral-NR80 as the seen (TR and VL) set and Archi-Stral-NR80 as the unseen (TS) set, several SeqFold2D models of sizes 400K-1.4M were trained for this cross-cluster study. As both intra- and inter-family sequence distributions are drastically changed by the de-redundancy procedure, model performances exhibit broad changes at the ensemble and family-specific levels. On the whole, all SeqFold2D models yield noticeably lower F1 scores on both seen and unseen datasets compared with the cross-sequence study, which can be attributed to the fact that the removed sequences are redundant and preferentially well-fitted. For each RNA family, its performance gain/loss is generally correlated with the increase/decrease of its population share in the dataset, consistent with the effects of observational bias on DL model development.

However, this cross-cluster study witnesses the emergence of significant TR-VL variances (i.e., overfitting) and further increases of the TR-TS variances (i.e., degraded generalizability) at both ensemble and family-specific levels for all SeqFold2D models, as shown in Suppl. Figs. S12&13. The TR-VL and TR-TS variances can be reduced with stronger regularization methods (e.g., larger dropout or weight decay rates). We carried out limited explorations but observed poorer overall performances for all datasets. Hence the SeqFold2D models presented here were developed with the same model hyperparameters as in the cross-sequence study. Rather curiously, the TR-VL variances typically appear when the F1 scores reach ~0.8 and further training improves the F1 scores for both datasets, but at a faster rate for the TR set. It is thus suggested that, in the early training phase, the directions with the deepest gradients are given by the sequence-structure correlations common to the TR and VL sets. The gradients become shallower as the training progresses and the models start fitting spurious patterns in the TR set while continuing to improve on the VL set. Consequently, given the more dissimilar sequence distributions between Stral-NR80 and Archi-Stral-NR80 than between the TR and VL sets taken from the same Stral-NR80 dataset, even larger TR-TS variances are observed for all SeqFold2D modes (Suppl. Fig. S11). Fig. 4 shows the TR-TS variances for the SeqFold2D-1.4M model for individual families, evidencing substantial generalization gaps for all families existing in both datasets. Overall, despite the excellent performances on the TR and VL sets (with the best F1 scores of 0.956 and 0.905, respectively, see Suppl. Fig. S12), the performances on the TS set are rather unsatisfactory on the absolute scale (the best F1 score of 0.762) and some RNA families have F1 scores close to or far below 0.5 (e.g., 16S and 23S rRNAs). These suggest that that the uses of non-redundant RNA sequences (below 80% herein) do not improve the generalizability of *de novo* DL models which however exhibit strong propensities towards overfitting.

**Figure 4.**
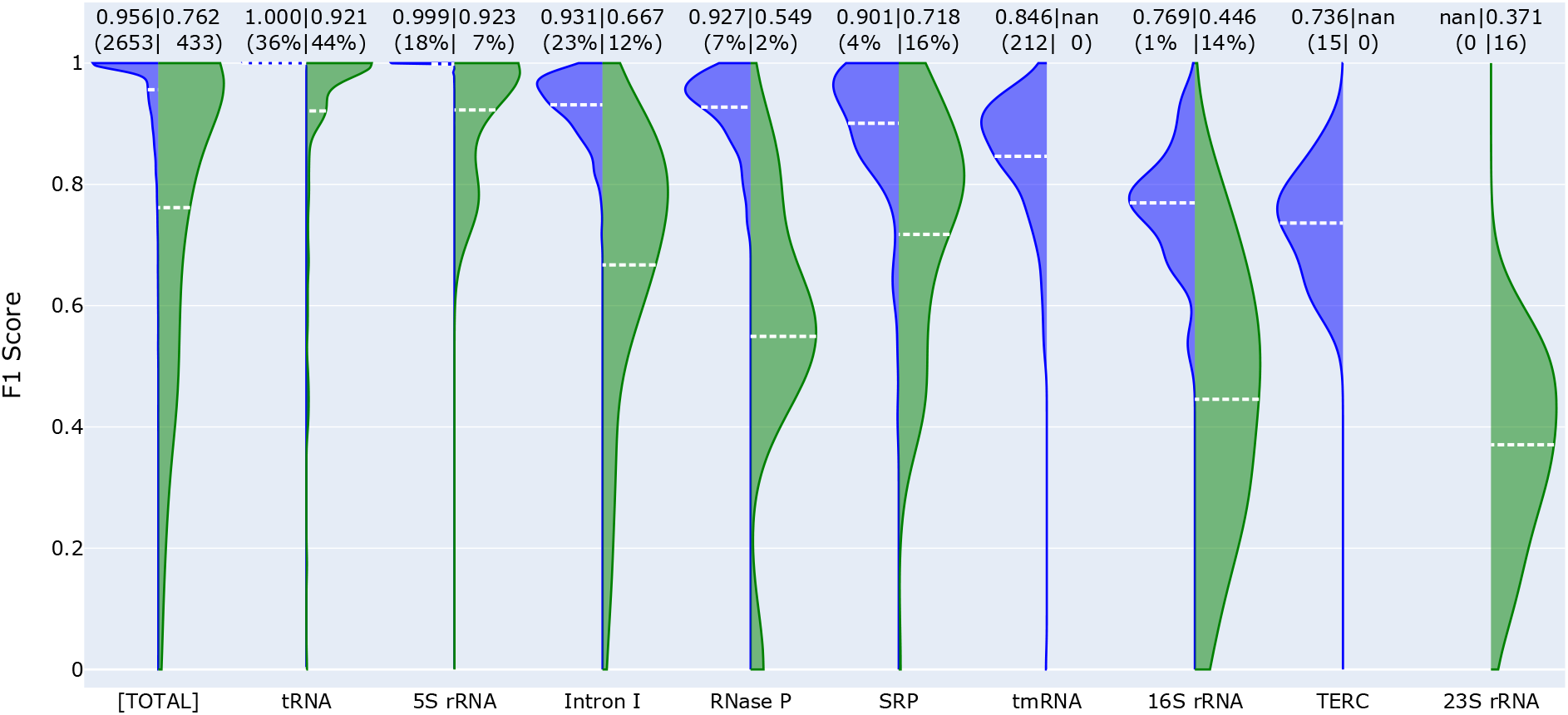
The F1 scores on the TR set from Stralign NR80-L600 (left, blue) and the TS set Archi-Stral-NR80 (right, green) by the SeqFold2D-1.4M model. The specific annotations follow that in Fig. 3. Note that tmRNA and TERC are absent in Archi-Stral-NR80 due to their high redundancies between the Stralign and ArchiveII datasets and that 23S rRNA only exists in the Archi-Stral-NR80 dataset.

To verify that the observations are not due to the relatively smaller model sizes or other insufficiencies of the SeqFold2D architecture, we trained several SeqFold2D models on the bpRNA TR0 and VL0 datasets and tested them with the bpRNA TS0 datasets. The three bpRNA sets (TR0, VL0, and TS0, a total of 13,419 structures) are all below 80% sequence similarity filtered with CD-HIT and much larger than the Stral-NR80 dataset (3,122 structures). When benchmarked against other models (Suppl. Fig. S14), the smallest SeqFold2D model with 960K parameters achieves an F1 score of 0.635 on the bpRNA TS0 dataset, only inferior to the Ufold-8.6M* model with F1~0.654. Larger SeqFold2D models generally lead to better performances, though with only marginal gains for the VL0 and TS0 datasets. Still, the largest SeqFold2D-3.5M model (N=7 and C=96) attains an F1 score of 0.665 on the bpRNA TS0 set, the highest reported to date. Notably, despite the relatively poor scores on the TS0 set, rather high F1 scores are observed on bpRNA TR0 with over 10K non-redundant sequences, 0.812 and 0.903 by SeqFold2D-960K and SeqFold2D-3.5M, respectively. This again testifies to the remarkable expressive capacity of DL models. However, even greater levels of TR-VL and TR-TS variances are also observed with the bpRNA datasets whose sequence distributions are expected to be more diverse. Similar to the model training with Stral-NR80, the TR-VL variances with bpRNA only appear in later training phases, though with an onset at a much smaller F1 value of ~0.55. Continued training increases the TR-VL variances but also the performances over the VL set (e.g., the best F1 scores of 0.657 and 0.686 for SeqFold2D-960K and SeqFold2D-3.5M, respectively). Importantly, the experimentation with the bpRNA dataset corroborates the challenges of *de novo* DL models in generalizing over non-redundant sequences even when developed with a large number of non-redundant sequences.

### Pairwise sequence alignment: decisive roles of sequence similarity in model generalization

Our observations so far indicate that both model training and test performances strongly depend on the sequence similarities within and between the seen and unseen datasets.In the case of high similarity as in the cross-sequence study, none or very little performance gaps can be achieved for non-identical sequences. On the other hand, the low sequence similarity in the cross-cluster study results in large overfitting and generalization gaps. Here we aim to probe the correlation between model generalization and sequence similarity quantitatively. With the model performance quantified by the F1 score, we carried out pairwise sequence alignment (PSA) as the means to assess the similarity between a test sequence and the seen dataset, as described below. The SeqFold2D-1.4M model with Stral-NR80 as the seen and Archi-Stral-NR80 as the unseen datasets is analyzed here because of their relatively small numbers of sequences with broad distributions of F1 scores.

For a given test sequence, it is first aligned against every sequence in the seen set (Stral-NR80, 3122 sequences in total) to produce 3122 PSAs. A percentage sequence identity (PSI) score is then calculated from each PSA as the number of identical aligning nucleotides divided by the average length of the sequence pair. Note that we experimented with alternative PSI definitions (e.g., using the shorter length of the sequence pair) and observed qualitatively consistent behaviors. As the seen set consists of highly diverse sequences across eight RNA families, the 3122 PSI scores are broadly distributed. Now with the pairwise PSI values of the test sequence to the entire seen set, the question we next ask is, can we use the subset of seen sequences above a certain PSI value to inform the model performance for the test sequence? One extreme case would be setting the PSI threshold to 1.0 (i.e., using a seen sequence identical to the test sequence), where the same performances are granted. Lowering the PSI threshold gradually includes more dissimilar sequences from the seen set and the informative power is expected to decrease. As such, the dependence of the informative power on the PSI threshold may provide quantitative insights into the model generalizability. Specifically, for the subset of seen sequences above a given PSI threshold, we average their F1 scores weighted by the PSI values to obtain the F1-seen score as the surrogate of their informative power. The F1-seen score is then compared with the actual F1 score for the test sequence (F1-unseen) at different PSI thresholds.

The entire process described above is repeated for all test sequences in the Archi-Stral-NR80 dataset (433 in total). As a result, at each PSI threshold, we obtain 433 pairs of F1-seen and F1-unseen values and examine their statistical correlations with rubrics such as the Pearson correlation coefficient (PCC). In order to rule out coincidental statistics, three different PSA programs were used: Foldalign (46), LaRA2 (47), and LocaRNA (48). Figs. 5A&B show the ratios and PCC values between the F1-unseen and F1-seen scores as a function of the PSI threshold. Consistent trends are given by the three PSA programs, supporting the robustness of observed dependencies. It is worth noting that all three programs find seen sequences with PSI values higher than 0.8, somewhat unexpected because of the below 80% redundancy between Stral-NR80 and Archi-Stral-NR80 filtered by CD-HIT. We speculate that this is caused by algorithmic differences as CD-HIT employs a greedy incremental clustering method that first estimates sequence similarity via short-world counting (word length of five used in this study). This may lead to missed matches by CD-HIT as the PSA programs always carry out actual sequence alignment. It is thus suggested that additional de-redundancy steps (e.g., with BlastN (49), Infernal (50)) may be warranted for situations where sequence non-redundancies are critical. Our analysis here is not significantly affected by low levels of highly similar sequences and it is reassuring that the F1 ratios and PCC values both approach the asymptotic value of 1.0 as the PSI threshold approaches 1.0. Thus no further de-redundancy was carried out.

**Figure 5.**
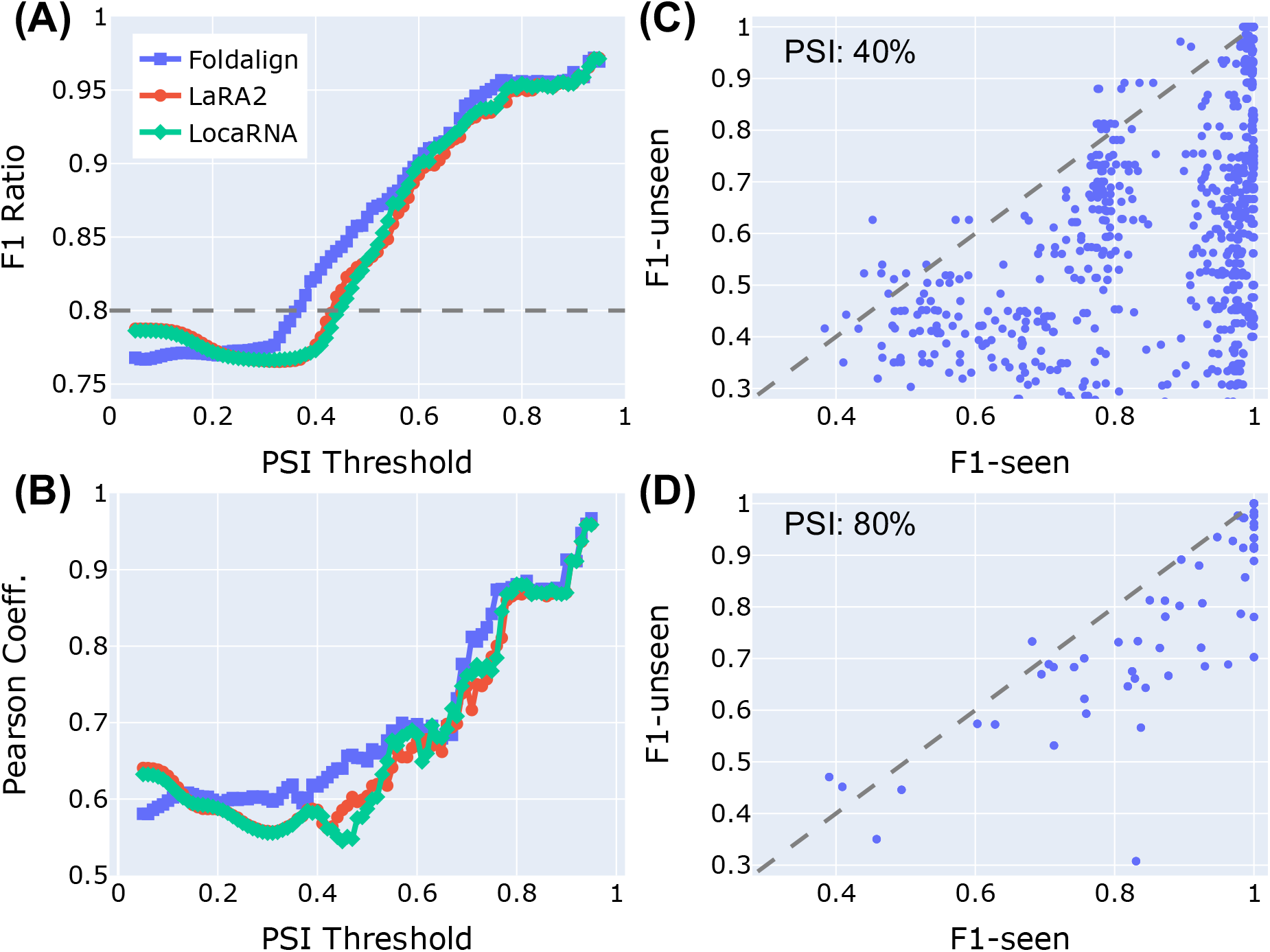
Illustrations of the correlations between the F1-unseen and F1-seen scores. (A) The F1-unseen over F1-seen ratios as a function the PSI thresholds. The horizontal dashed line marks the F1 ratio between the entire unseen and seen datasets. (B) The PCC values as a function of the PSI thresholds. (C) The distributions of the F1-unseen and F1-seen scores at the nominal PSI threshold of 40% by Foldalign. (D) The distributions of the F1-unseen and F1-seen scores at the nominal PSI threshold of 80% by Foldalign. Note that it is common to find no seen sequences for a test sequence above a high PSI threshold, leading to many null F1-seen values that are absent in (D).

The main observation from Fig. 5 is the rapid declines of both F1 ratios and PCC values with the decrease of sequence similarity. In the direction of decreasing PSI thresholds, both F1 ratios and PCCs start out in the high 90%s, affirming the excellent generalizability over nonidentical but similar sequences observed in the cross-sequence study. The F1 ratios and PCCs then quickly drop as the PSI threshold is lowered, indicating a fast decay of generalizability over increasingly dissimilar sequences. The F1 ratio approaches its ensemble value (i.e., the F1 ratio between the entire unseen and seen datasets) around PSI ~ 0.4, in spite of the existence of significant PCC values of ~0.6 for all curves. Representative distributions of F1-seen and F1-unseen values at two PSI thresholds are shown in Figs. 5C&D. Overall, considering both F1 ratios and PCC values, the informative power of the seen set falls below PCC ~ 0.8 at PSI ~ 0.7 and diminishes completely at PSI ~ 0.4. While the exact PSI transition points likely depend on the specific datasets used, the PSI definition, etc., our observations provide quantitative insights on how the generalizability of *de novo* DL models depends on the sequence distributions of the seen and unseen datasets.

### Cross-family study: inability to generalize over unseen RNA families

An even more stringent examination of model generalizability is to train DL models with one RNA family and test with other types. While all *de novo* DL models are expected to fail such a cross-family test, we experimented with a single SeqFold2D-400K model trained on the most abundant family type, tRNA. To maximize the number of tRNA sequences, the Stralign and ArchiveII datasets were combined into one Strive dataset with a total of nine families (see Suppl. Fig. S5). Training was relatively straightforward given the high similarity levels of the tRNA sequences. To limit overfitting, we stopped training when the TR-VL variance became significant around the F1 score of 0.975. As shown in Suppl. Fig. S15, the SeqFold2D-400K model fails completely on all other RNA families, yielding the F1 scores ranging from 0.03 to 0.1. We did not pursue the use of other families for training in part due to their relatively smaller populations. We like to note a recent proposal of a different cross-family test by only excluding the test family type from the seen set (35). It was shown that *de novo* DL models performed poorly on such cross-family tests, consistent with our observations.

## CONCLUSION AND DISCUSSIONS

In this study, we set out to investigate the expressive capacity and generalizability of *de novo* DL models by varying their sizes and the sequence distributions of the seen and unseen datasets. To this end, we design the SeqFold2D architecture with a minimal two-module structure and without any post-processing steps such as model averaging and grid search. The SeqFold2D models demonstrate remarkable learning power of the relationships between RNA sequences and secondary structures by achieving excellent performances for all training sets, e.g., the SeqFold2D-960K model attains the F1 scores of 0.985, 0.971, and 0.812 for the Stral-ND, Stral-NR80, and bpRNA TR0 sets, respectively. This is especially notable for the bpRNA TR0 dataset comprising over 10K non-redundant sequences. For the unseen/test sequences, the SeqFold2D models outperform all other *de novo* DL models and traditional algorithms, despite often with much fewer parameters and without post-processing. However, while the SeqFold2D models are able to generalize well over unseen sequences at the cross-sequence level, significant performance gaps appear at the cross-cluster level, manifested as sizeable TR-VL variances (overfitting) and even larger TR-TS variances (poor generalizability). With the help of pairwise sequence alignment, we quantitatively analyze how model generalizability depends on the sequence similarity between the seen and unseen sequences and reveal a rapid decline in performance for increasingly dissimilar sequences. Altogether, these observations suggest that the class of *de novo* DL models are largely statistical learners of RNA sequence and structure pattern mappings and that their applicability in the real world is determined by the sequence distributions of the seen and unseen/test datasets.

Meanwhile, our study can be extended in several important directions. First, the SeqFold2D models were manually tuned to maximize the performances on the VL set rather than to minimize TR-VL or TR-TS variances (e.g., via early stopping or stronger regularization). We found this strategy also led to best performances on the TS set, which can be argued to be optimal for the purpose of benchmarking. In essence, the models are fitting both general and spurious patterns at the same time in later training phases and hence certain levels of overfitting are beneficial to test performance. Nonetheless, model hyperparameters can be further explored to better balance performances and generalizability. Second, we primarily focused on the F1 score as the sole metric for model performance and have yet to analyze other metrics such as Precision, Recall, pseudo-knot prediction, and long-range contact prediction that have been discussed by other DL models. We also focused on large datasets only and have not trained SeqFold2D models on much smaller datasets derived exclusively from PDB structures. These additional studies are work in progress. Lastly, it would be highly desirable to understand and explain how the *de novo* DL models predict secondary structures. Existing DL models encompass a wide range of network architectures, e.g., U-net by Ufold, 2D-LSTM by SPOT-RNA, and 1D-LSTM by MXfold2. We experimented with both 1D-LSTM and selfattention-based transformer layers and found similar performances. Various techniques from the machine learning field can be used to inspect the hidden 1D and 2D representations and probe the inner workings of these diverse DL models (51). For example, model interpretability may help us understand why a SeqFold2D model of the same size would overfit the larger bpRNA TR0 more than the Stralign NR80 dataset.

Furthermore, continued developments of the class of *de novo* DL models, holding the advantage of requiring no more information other than nucleotide sequences, are much needed to improve both performance and generalizability. For example, with or without postprocessing, no DL models have accomplished satisfactory performances on the bpRNA TS0 set, with the highest F1 ~ 0.665 by the SeqFold2D-3.5M model in spite of an F1 score of 0.903 on the bpRNA TR0 set. The causes of such poor generalization, as suggested by this study, are likely rooted in the statistical learning nature of DL models. As such, the ideal solution is to increase the number and distribution of training structures. Direct experimental determinations of RNA secondary structures are however slow and costly (52) and high-throughput measurements such as PARS (53) and SHAPE (54) usually yield nucleotide-level activity profiles rather than base-pairing partners. Note that high-throughput experimental data can serve as valuable constraints for structure prediction (55,56). Given that comparative sequence analysis is the main method of data curation for large structure databases, renewed efforts with more sophisticated pipelines such as RNAcmap (57) and rMSA (https://github.com/kad-ecoli/rMSA2) could also be viable. On the other hand, before larger structure databases become available, wide ranges of resources and techniques in the fields of biology and machine learning may be exploited to develop accurate and generalizable *de novo* DL models, some of which are discussed below.

> *Input enrichment*. Input information may be enriched beyond one-hot identifiers of RNA sequences. Chemical and physical properties of the nucleotides can be embedded to capture their intrinsic features particularly about the different bases, though one potential caveat is that nucleotides are much less diverse compared with amino acids. Moreover, unsupervised pre-training of known RNA sequences with natural language models such as BERT (58) may offer a powerful pathway for representation learning of RNA sequences. One example of this kind is the recent DNABERT model pre-trained on human genome sequences (59). In the case of RNA, non-coding RNA sequences in the RNAcentral database (60) can be used for pre-training such that richer representations can be learned from all non-coding RNAs.
>
> *Architecture design*. Avenues can be explored to infuse the mechanisms of RNA structure and folding (3) into the DL architecture. At the sequence and pair-interaction representations, one remarkable example is the EvoFormer module in the Alphafold2 architecture (61) that facilitates cross-representation exchanges and builds in inductive biases such as triangular updates. A very recent work, DeepFoldRNA, demonstrated that an EvoFormer-like architecture also works well for RNA (62). This however requires the use of multiple-sequence alignment and it remains to be seen whether good effects can be obtained without co-evolutionary information. Alternatively, it may be possible to introduce learning biases in the form of auxiliary loss functions similar to the Physics-Informed Neural Networks (PINN) (63). For example, the structural constraint of no base multiples can be enforced by a term 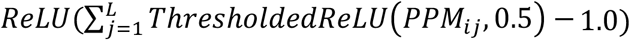 and sharp turns can be penalized through a masked loss of the diagonal elements. Another important aspect of RNA secondary structures is their topology (64). A *posteriori* method is to increase the weights of loop opening and closing base pairs in the loss function.
>
> *Multi-task learning*. Model outputs can also go beyond base-pairing matrices. Additional structural properties can be predicted at the nucleotide level, such as nucleotide-wise solvent accessibility or activity profiles that can be measured independently (e.g., via SHAPE). Thermodynamic quantities at the sequence level such as free energies and melting temperatures may also be predicted. Perhaps one most practical model prediction would be the confidence levels of the predicted secondary structures, like the pTMscore by Alphafold2. We first exploited the fact that each predicted PPMij is a probability itself and has an apparent variance of *PPM_ij_* (1 —*PPM_ij_*) under the assumption of a binomial distribution. One can derive the variance of the resultant soft F1 score which is function of the predicted PPM and the ground truth. Suppl. Fig. S16 shows the correlations between the estimated variances and the values of F1 scores for the SeqFold2D-960K model developed with the bpRNA TR0 and VL0 datasets. While clear negative correlations are observed when the F1 scores are greater than 0.8, the estimated variances show no or even reversed correlations when the F1 scores fall below 0.8, rendering it completely uninformative of model confidence. Additionally, we experimented with the addition of a third module to predict the F1 score which however showed poor generalization power despite excellent performances on the seen sequences. More work is needed to overcome the barrier of model generalizability.

In closing, RNA secondary structures confer important biochemical and physical features to all RNAs, coding or non-coding, and they often play critical roles in the biological functions of RNA molecules. The challenges of predicting RNA secondary structures *de novo* have inspired the development of numerous DL models that demonstrate unprecedented expressive power as well as stern reliance on broadly distributed training data. This study provides quantitative analyses of the model capacity and generalizability in the context of different sequence distributions. And various pathways for future advances are discussed so as to catalyze the development of next-generation *de novo* DL models for RNA secondary structure prediction.

## ACKNOWLEDGEMENT

We thank Dr. Chengxin Zhang for the suggestion of LaRA2 for pairwise sequence alignment and the GW High Performance Computing Facility for the free access of CPU and GPU nodes.

## Notes

### Competing Interest Statement

The authors have declared no competing interest.

### Summary of Updates

The major changes are in the last few paragraphs of the Conclusion and Discussions section. Some important citations were added for completeness. We did minor polishing throughout the maintext and supplemental materials. No changes were made on figures.

