## Supplementary material for "Decisive Roles of Sequence Distributions in the Generalizability of *de novo* Deep Learning Models for RNA Secondary Structure Prediction": Suppl. Materials and Figures

Xiangyun Qiu

Department of Physics, George Washington University, Washington DC 20052

### MATERIALS AND METHODS

#### Datasets

*The RNA Stralign dataset.* The RNA Stralign dataset was curated in 2017 (1) for RNA sequence alignment and secondary structure predictions (hence highly redundant). Being relatively modern, Stralign was noted to have greater sequence diversity than previous datasets (e.g., BRAliBase (2)) and comprises several families longer than 320 nucleotides (up to 1851). The Stralign dataset has a total of 37,149 sequences distributed over eight RNA families as shown in Fig. S1. For the purpose of deep learning (DL) model development, duplicated sequences are removed and sequence lengths are usually limited to 600 nucleotides due to computational constraints, resulting in 20,118 sequences referred to as the Stralign ND-L600 dataset or Stral-ND in short. Overall, the sequence distributions in the Stral-ND dataset are highly uneven, presenting steep observational biases for training DL models. This is first reflected in the imbalanced representations of different RNA families, e.g., the top two families (5S rRNA and tRNA) account for nearly 80% and the bottom four families for less than 8%. Another is the family-specific, uneven sequence length distributions (shown in Fig. S2). It is noteworthy that the dataset contains many sub-domains taken out of full-length sequences.

As structure is more conserved than sequence, non-identical sequences can give highly resembling structures, adding another source of observational bias. One common mitigation is to remove similar sequences above certain 80% sequence identity level which has been shown to the inflection point of sequence-structure correlations (3). We consequently obtained such dataset, denoted as Stral-NR80, by reducing the Stral-ND dataset with the program CD-HIT to below 80% sequence identity (80% is also the lowest allowed by CD-HIT). As shown in Fig. 1A and Fig. S1, this led to a dramatic reduction in size for Stral-NR80, with just 3,122 sequences or ~1/7$^{th}$ of the Stral-ND dataset. The most populous families (16S and 5S rRNA and tRNA) all have high levels of redundancy as large as 20 folds, whereas the less-represented families typically show less than 3-fold redundancy. Out of curiosity, we also verified that all cross-family sequence pairs are below 80% identity level as expected. Therefore, in the context of the entire polynucleotide sequence space, these RNA families can be viewed as distinct clusters with inter-family dissimilarities (or distances) at least 80%, while each cluster itself also spans beyond 80% similarity levels. The exact intra- and inter-family dissimilarities are however unknown. As mentioned, each RNA family further has characteristic length distributions as shown in Fig. S2. These fundamental differences all present challenges for DL models to generalize over different families.

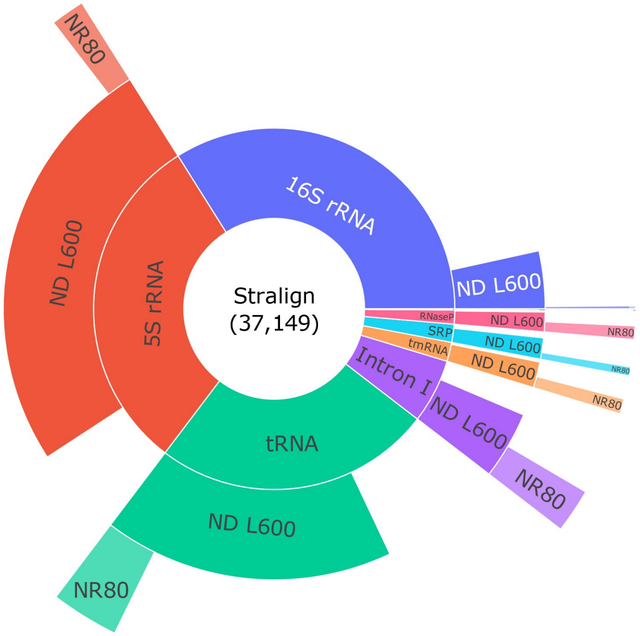


Figure S1. The population distributions of RNA families in the RNA Stralign dataset at different sequence redundancy levels. Note that this is an unscaled version of Fig. 1 in the main text and that the TERC (telomerase RNA) population is too small and barely visible. The innermost ring shows the distributions of the eight RNA families in the original Stralign dataset. The arches along the middle ring, labelled "ND L600", show the population of each RNA family in the Stralign ND-L600 dataset after removing duplicated sequences and sequences longer than 600 nucleotides. The outermost ring, labelled "NR80", shows the RNA family distributions in the Stralign NR80-L600 after removing redundant sequences above 80% sequence identity.

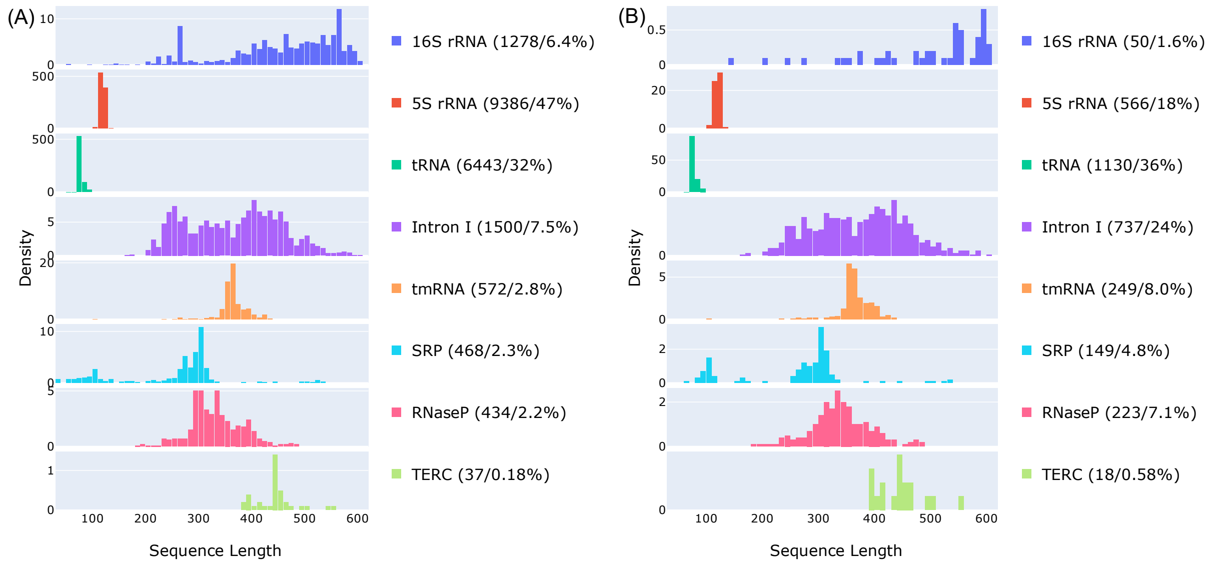


Figure S2. The length distribution of each RNA family in the Stralign ND-L600 (A) and Stralign-NR80-L600 (B) datasets shown in the same order and color as in Fig. S1. The number of each family type and its percentage in the parent dataset are shown in the legend.

*The RNA ArchiveII dataset.* The RNA ArchiveII dataset (4) is a collection of benchmarking RNA secondary structures determined by comparative sequence analysis. It is worth noting that the ArchiveII dataset has no non-canonical base pairs and no pseudoknots. Structures with unknown residues were also omitted and long sequences were divided into domains no longer than 700 nucleotides. In addition to the eight RNA families as in the Stralign dataset, ArchiveII includes two more families with longer sequences: 23S rRNA and group II intron, despite its much smaller size of 3,975 sequence in total (~11% of Stralign). As done for Stralign, we obtained ArchiveII ND-L600 dataset (Archi-ND, 3395 sequences) by removing duplicates and sequences longer than 600 nucleotides. Note that all group II intron sequences are longer than 600 and thus absent in the Archi-ND set. The Archi-NR80 set (1221 sequences) was obtained by removing redundant sequences above 80% sequence identity. The population distributions and length distributions of all RNA families are shown in Suppl. Figs. S3 and S4. For the cross-cluster study, we further removed the sequences in the Archi-NR80 dataset with above 80% sequence similarity level with the Stral-NR80 dataset, yielding the Archi-Stral-NR80 dataset (433 sequences).

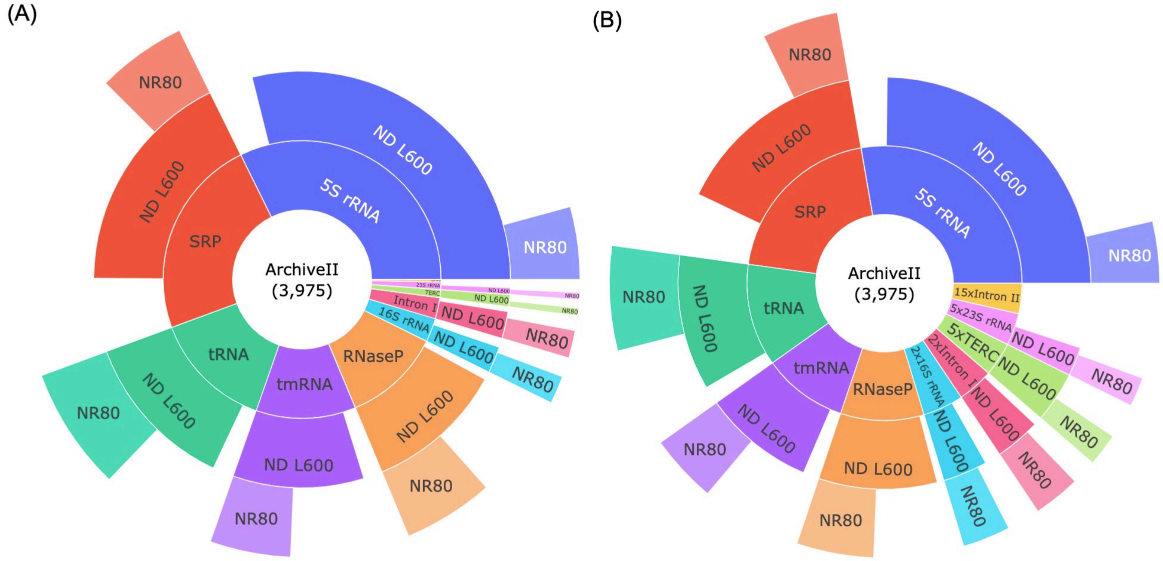


Figure S3. The population distributions of all RNA families of the RNA ArchiveII dataset at different sequence redundancy levels. Two versions for the same underlying datasets are shown, the unscaled version (A) and the scaled version (B) for visibility of the underrepresented families that are scaled up by the multiplier N shown in the label. With the same notations as used in Fig. S1, the innermost ring shows the relative populations of the RNA families in the original ArchiveII dataset. The arches along the middle ring show the population distributions of the RNA families in the ArchiveII ND-L600 dataset after removing duplicated sequences and sequences longer than 600 nucleotides. The outermost ring shows the distributions of the RNA families in the ArchiveII NR80-L600 after removing redundant sequences above 80% sequence identity. Note that group II intron, labelled as Intron II, all have lengths longer than 600 and are thus absent in the ND-L600 and NR800-L600 datasets.

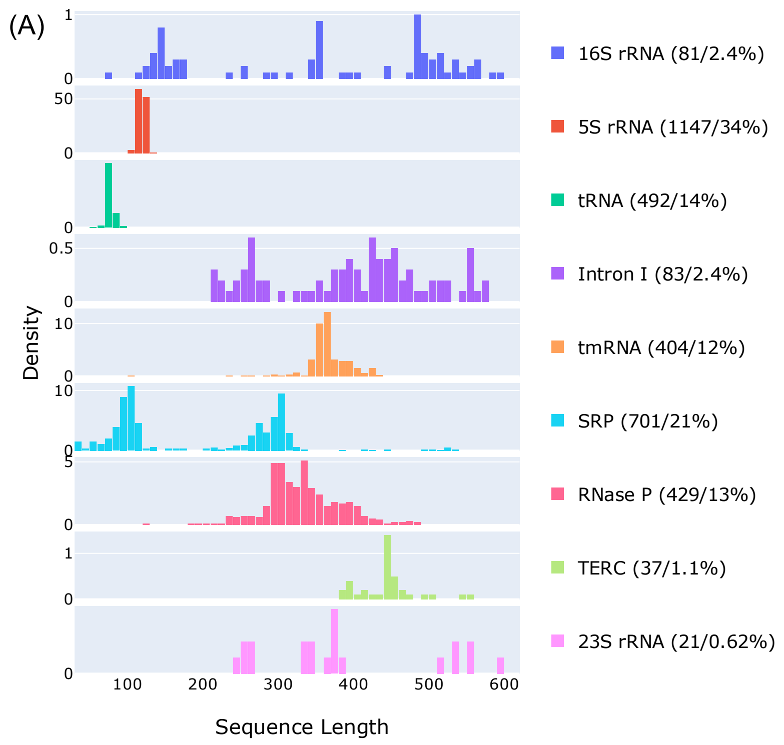


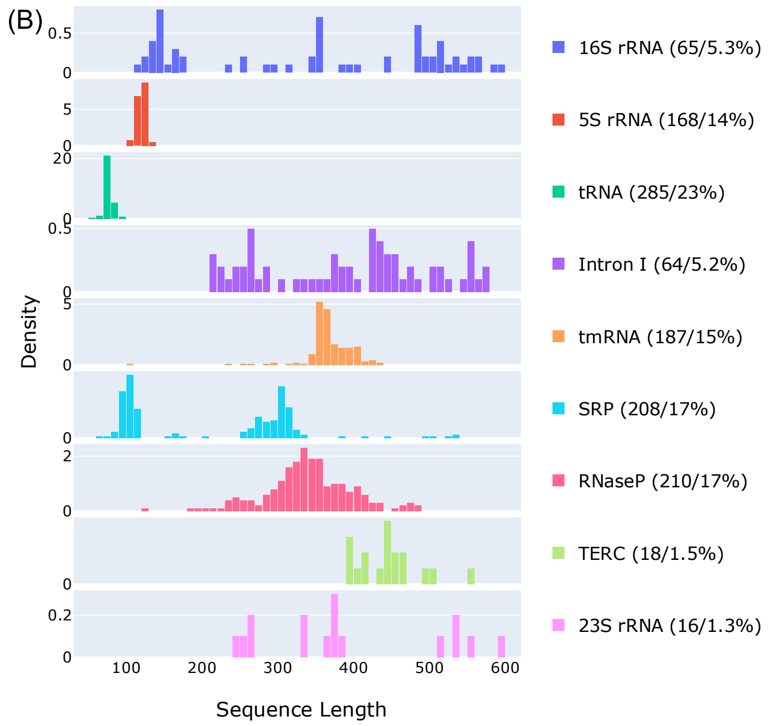


Figure S4. The length distributions of all RNA families in the ArchiveII ND-L600 (A) and NR80-L600 (B) datasets. The order of the RNA families shown follows that of the Straligin datasets in Fig. S2 to facilitate comparison, rather than by the order of population as in Fig. S3.

*The Strive dataset.* The Strive dataset is the sum of the RNA Stralign and ArchiveII datasets with duplicated sequences removed. It is compiled mainly for the cross-family study. We followed the same procedure as done for the Stralign and ArchiveII datasets to obtain the Strive ND-L600 and Strive NR80-L600 datasets, the RNA family distributions of which are shown in Fig. S5. Specifically, the Strive ND-L600 contains non-duplicate sequences up to 600 nucleotides and the Strive NR80-L600 further removes sequences above 80% similarity levels with CD-HIT. Fig. S5 shows the distributions of RNA families in the Strive ND-L600 and NR80-L600 datasets.

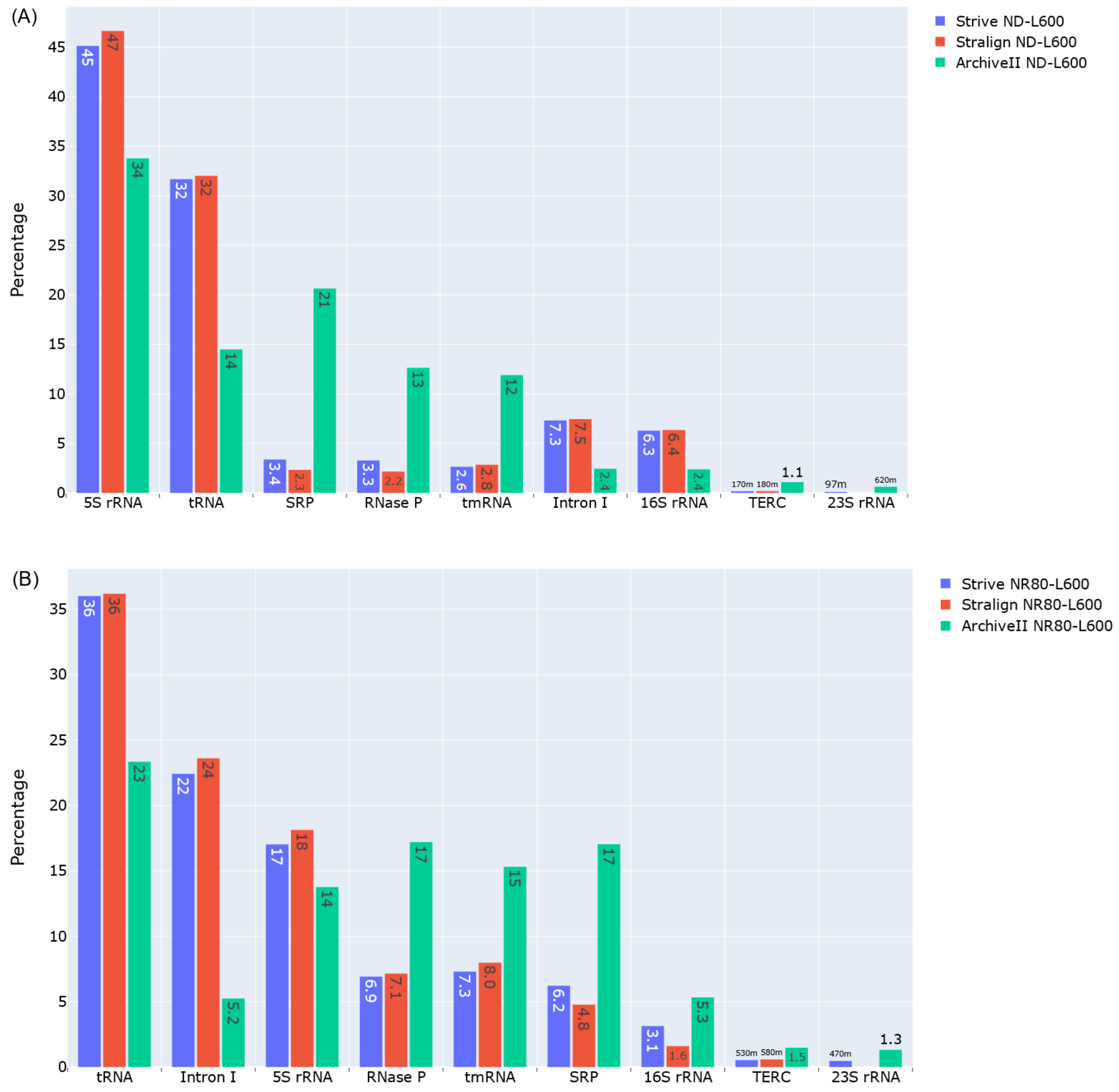


Figure S5. The distributions of RNA families in the Strive ND-L600 dataset (A) and the Strive NR80-L600 dataset. Together shown are the corresponding datasets of Stralign and ArchiveII for comparison. The order of RNA families is sorted by the family abundance in the Strive datasets.

*The bpRNA dataset.* The bpRNA-1m dataset (5) is a >100K RNA secondary structure collection (102,348) from seven databases: the comparative RNA web (CRW) site (55,600), tmRNA (728), SRP (959), SPR (623), RNP (466), RFAM (43,273), and PDB (669). Despite being the largest database for RNA secondary structures, its sequence distributions are highly uneven across its member databases. Like its member databases, the vast majority of the secondary structures in the bpRNA dataset are determined by comparative sequence analysis. The downloadable form however only provides the source (e.g., CRW or SRP) rather than the actual RNA family type. We thus chose Stralign and ArchiveII over bpRNA for detailed studies. For the development of DL models, one commonly used bpRNA-derived dataset is the TrainSet0 (TR0, 10, 814 sequences), ValidSet0 (VL0, 1300), and TestSet0 (TS0, 1305) compiled by the SPOT-RNA team (6), with a total of 13,419 sequences. Specifically, sequence lengths are limited to be within 30 and 500 nucleotides, the sequence identity is trimmed to 80% with CD-HIT, and all sequences with high similarity levels to the PDB datasets are removed. Fig. S6 shows the source and length distributions of the bpRNA TR0 and VL0 sets. The same dataset choices (i.e., TR0, VL0, TS0) were used by Ufold and MXfold2 with pre-trained parameters available, noting that MXfold2 further removed non-canonical base pairs and pseudoknots from the RNA structures.

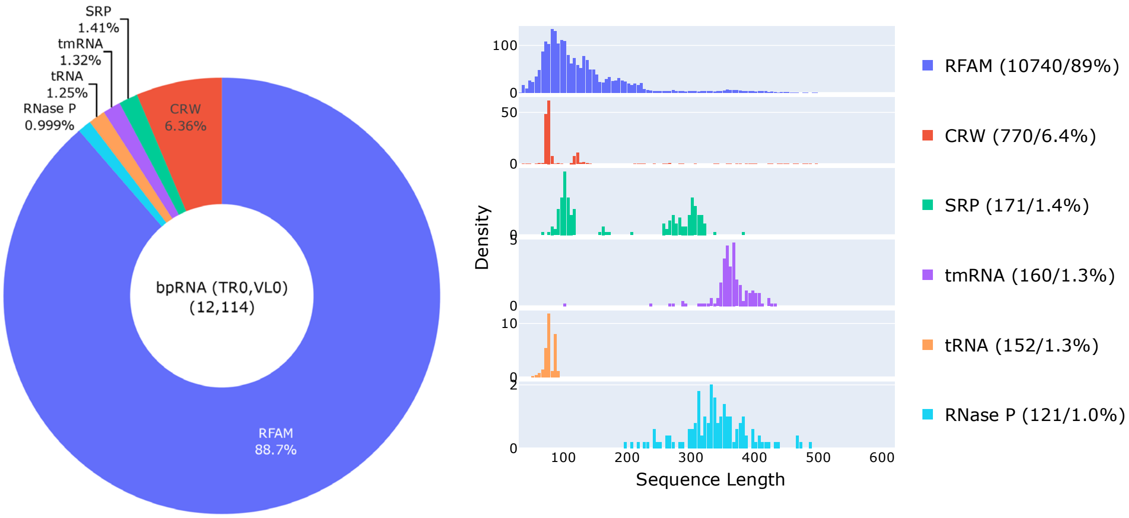


Figure S6. The population distributions (left panel) and length distributions (right panel) of the sequences grouped by their sources in the bpRNA TR0 and VL0 datasets. The x-axis range is shown up to 600 nucleotides for easy comparisons with Figs. S2&4, while the actual sequences are all shorter than 500 nucleotides. The TS0 set has essentially the same source and length distributions and thus not shown separately.

*The bpRNA-NEW dataset.* The bpRNA-NEW dataset was compiled by the MXfold2 team (7)and it is based on the newly added ~1500 families to Rfam 14.2 since Rfam 12.2 used by the bpRNA-1m dataset. It has a total of 5401 sequences with lengths shorter than 500 nucleotides and sequence identities below 80% filtered by CD-HIT. However, both non-canonical base pairs and pseudoknots are removed from the database. The level of base pairing is relatively low with an average of ~45%. These secondary structures are more likely underestimates of the true levels of base pairing, particularly for these families without 3D RNA structures providing seeding secondary structures. Fig. S7 shows the length distribution of the bpRNA-NEW dataset.

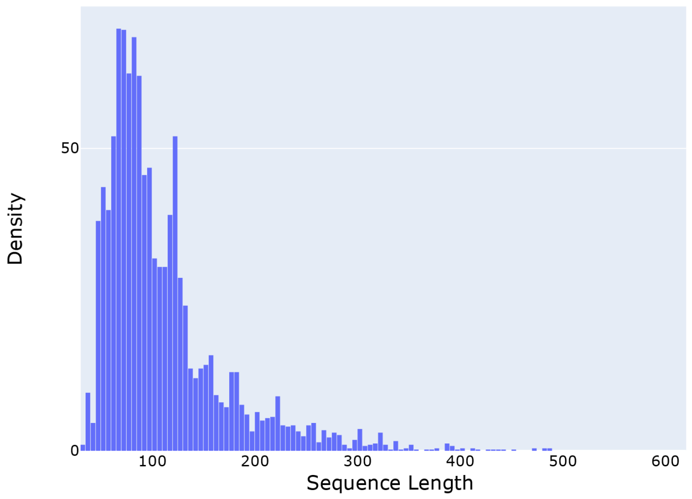


Figure S7. The length distribution of the bpRNA-NEW dataset. It is qualitatively similar to the Rfam length distributions of the bpRNA TR0 and VL0 sets shown in Fig. S6.

**Network architecture and training procedure**

With 1D RNA sequences as inputs and 2D pairing probabilities as outputs, the overall network architecture has two main learning modules, as shown in Fig. 1A in the main text. The first module consists of N1 stacked blocks of bidirectional Long-Short-Term-Memory (LSTM) or self-attention-based transformer encoders to learn richer 1D sequence representations, which are then transformed into 2D pair representations via outer-product. The second module consists of N2 stacked blocks of residual 2D convolutional layers to infer inter-nucleotide interactions. To reduce the design space of model hyperparameters, the numbers of blocks in both modules are kept the same, i.e., N1=N2=N. Layer normalization and dropout (0.2-0.42) layers are always applied after multiplications and additions with trainable weights and biases (except for the final output layer), where non-linear activations (LeakyReLU or Swish) are applied before. Note that the name Seqfold has been used for a method for reconstructing RNA structures from high-throughput sequencing data (8) and for another program (http://github.com/Lattice-Automation/seqfold), while we name our architecture SeqFold2D to emphasize the use of sequences as the only inputs and the output of 2D PPMs. Detailed description of each component is given below.

*Sequence embedding*. One-hot vectorization is used to digitize each nucleotide as a 1×4 vector, e.g., A as [1,0,0,0], C as [0,1,0,0], and an all-zero vector Z ([0,0,0,0]) for padding sequences to the same length. We further adopt a k-mer (k=3) representation for each base to include its neighbors. For example, ACGU is represented as four tokens of ZAC, ACG, CGU, and GUZ. Each RNA sequence of length L starts as a vector of shape L×12, which then passes through one feed-forward layer to obtain its embedding vector of shape L×C, where C is the channel size as a model hyperparameter. We further keep the channel size C the same throughout the model. As a result, the model size in terms of the number of parameters is largely determined by two design variables, N and C.

*Input block.* Two feed-forward layers are used to further mix the different channels while keeping the tensor shape as L×C. It can be argued that these feed-forward layers are not absolutely necessary, though no ablation studies were conducted.

*Module 1: 1D sequence encoding.* Each block, repeated N times in the module comprises one LSTM or transformer encoder layer. In the case of LSTM blocks, normalization and dropout layers are added between blocks and no additional activation layers are used. In the case of transformer encoders, sinusoidal positional embedding is added before the first block. A constant head size of 16 is used for the multi-head self-attention, under the condition that the channel size C is a multiple of 16. Similar performances are observed with the use of either LSTM or transformer encoders for this module.

*1D to 2D transformation.* For the transformation from the L×C 1D representation to 2D L×L×C pair representation, we experimented with outer-concatenation and outer-product and found similar performances. Outer-product is the usual choice to maintain the same channel size.

*Module 2: residual 2D convolution.* Each residual block comprises two 2D convolutional layers with kernel sizes of 5×5 and 3×3, respectively. The residual connection is done after activation and normalization layers to maintain a straight path for the so-called skip connection. It operates on the pair representation (L×L×C) and aims to facilitate the communication of each specific pairs with neighboring pairs.

*Output block.* The output block comprises of three fully connected layers that operate across the channel dimension only, i.e., no more communication between neighboring pairs. The dimension of the final layer is L×L×2 and Softmax is applied to get the matrix of unpaired probabilities and pairing probabilities, with the latter used for loss and metrics calculations.

*Evaluation metrics.* Given the predicted pairing probability matrix (PPM) and the ground truth (or Label) for an RNA sequence of length L, the F1 score defined as $\frac{2\times Precision\times Recall}{Precison+Recall} = \frac{2\times TP}{2\times TP+FP+FN} = \frac{2\times TP}{L^2+TP-TN}$, where Precision is TP/(TP+FP), Recall TP/(TP+FN), TP the number of true positives, FP false positives, FN false negatives, and TN true negatives. To compute the F1 score, the continuous PPM is discretized as 0 or 1 with a threshold of 0.5 without grid search.

*Loss functions.* The F1 score defined above cannot be directly used as the loss function because discretization renders it undifferentiable. A common surrogate is to compute the cross-entropy (CE) or square-error (SE) loss between $PPM_{ij}$ and $Label_{ij}$ for every i-j pair before averaging, which shares the same global optimum as the F1 score. Notably, the lopsided distribution of negative labels (i.e., 0s) creates an effortless slope towards the local minimum of predicting all zeros for PPM in the early phase of training and a weight bias of 300 for positive labels was used by E2Efold and Ufold to restore the balance. We however found this weight bias or the use of focal loss (9) as done in the image classification to artificially increase the false positives and chose not to apply such weight biases. Instead, we adopt a soft F1 score function as the surrogate loss to directly optimize the F1 score. The soft F1 score is straightforward to implement and was also used by E2Efold, as it simply bypasses the PPM discretization when calculating TP, TN, FP, and FN values, which makes it differentiable.

*Staged training.* Typical training starts with the CE loss function till the F1 score for the validation set stops improving. Then, the loss function is switched to the soft F1 score. This two-stage procedure was found to give the best F1 scores compared with using only one type of loss function.

*Hyperparameter turning.* To limit the number of searches, we tuned one hyperparameter at a time while also taking into considerations of the best practices in the literature and by other models. The SeqFold2D models of different sizes were tuned separately and the number of epochs for tuning is usually limited to 150 total for efficiency. For example, we fixed dropout to 0.25 and weight decay to 0.01 when tuning the learning rate between 1e-2 and 1e-6. After finalizing the learning rate (usually between 1e-3 and 1e-4), we proceeded to tune dropout between 0.1 and 0.6 and found optimal dropouts to be between 0.2 and 0.42 (usually larger rates for larger models). We did not tune batch size which is set to be the largest allowed by the GPU memory (usually between 8 and 16). The rubric for the best model is based on the F1 score on the validation set.

**RESULTS**

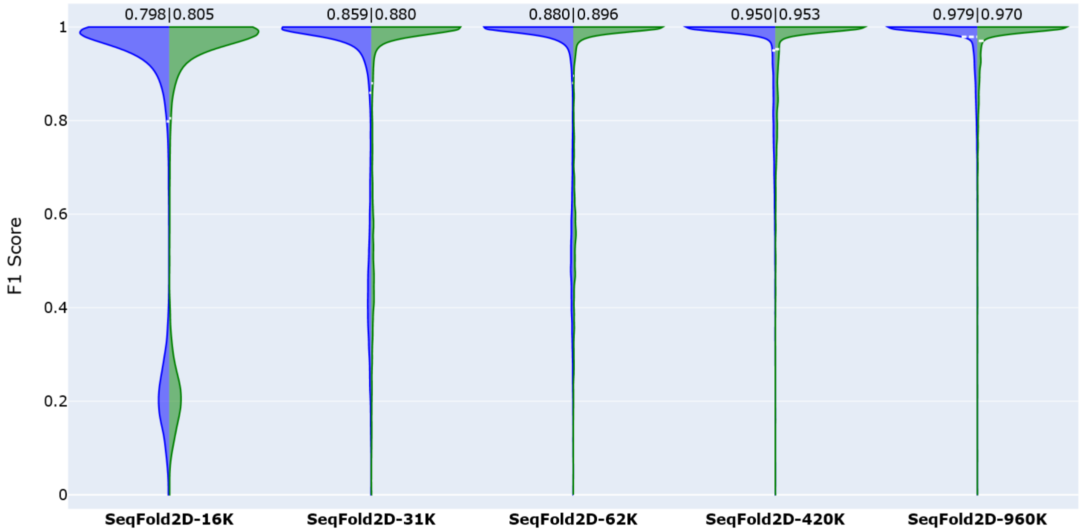


Figure S8. The F1 scores of the training (left, blue) and validation (right, green) sets for the SeqFold2D models developed with the Stralign ND-L600 (Stral-ND) dataset randomly split into three subsets: training (TR), validation (VL), and test (TS). The averaged F1 scores are shown at the top and also as dashed lines (white) within the corresponding violin plots (often too narrow to be spotted). Very little TR-VL variances are observed, indicating that the SeqFold2D models are describing the distribution of the entire Stral-ND dataset while being trained on the TR subset of the distribution. Note that the F1 scores were saved during training and all dropout layers were active for the TR set but not for the VL set. These make the F1 scores shown here slightly lower than the values computed without dropout.

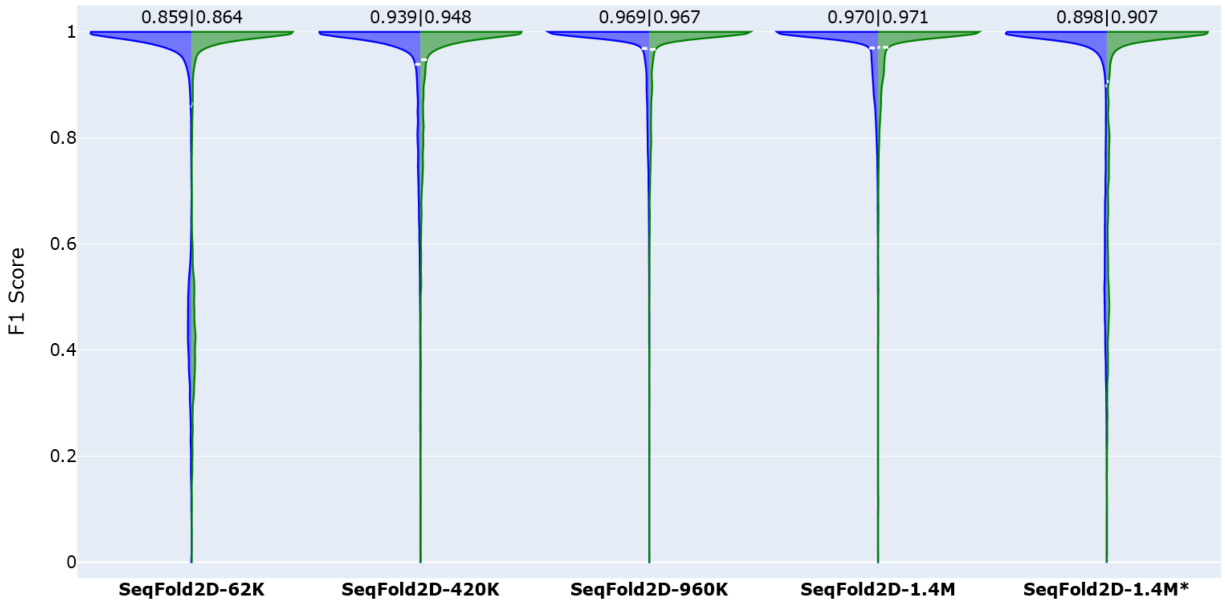


Figure S9. The F1 scores of the training (left, blue) and validation (right, green) sets for the SeqFold2D models developed with the Stralign ND-L600 (Stral-ND) dataset randomly split into two subsets only: training (TR) and validation (VL). The test set is the ArchiveII ND-L600 dataset as presented in the main text. The main difference between this set of SeqFold2D models and those in Fig. S8 (with Stral-ND split into the TR, VL, and TS sets) is the slightly larger TR set used here, while the training hyperparameters are kept the same for models with the same size. Somewhat surprisingly, this set of models show slightly lower F1 scores for the TR set compared with those shown in Fig. S7. We do not have good explanations for the drops and did not further investigate the causes as the F1 scores for the VL set are very close. The SeqFold2D-1.4M* model was trained following the similar choices made by E2Efold and Ufold, specially with the cross-entropy loss function only and a weight of 300 for positive labels. As the shown TR and VL F1 scores were saved during training without post-processing, the scores from the SeqFold2D-1.4M* model are significantly lower than that after post-processing. For example, the averaged F1 score for the TR set increases from 0.898 to 0.981 with post processing for SeqFold2D-1.4M*.

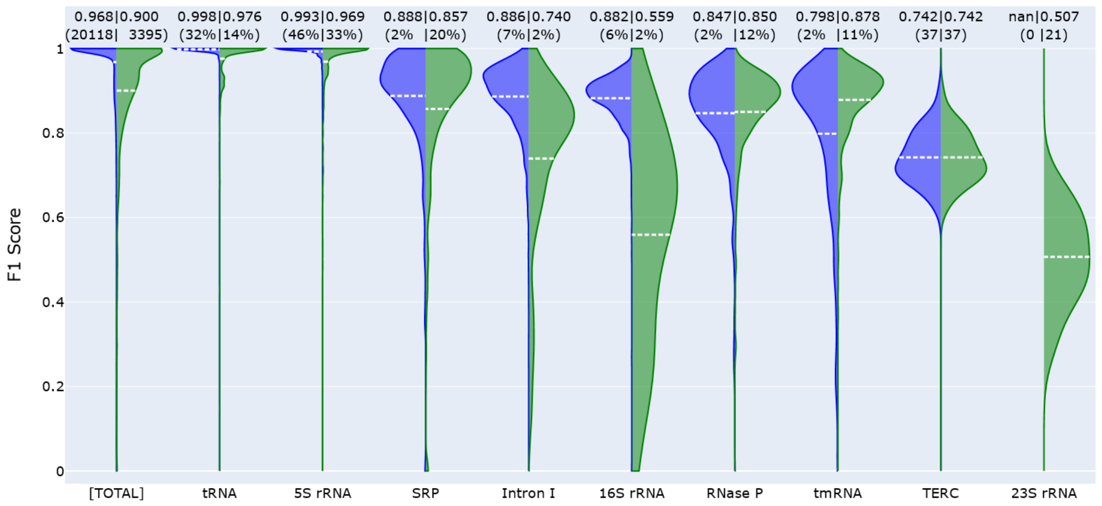


Figure S10. The F1 scores of the seen set (Stralign ND-L600, left in blue) and the unseen/test set (ArchiveII ND-L600, right in green) for the Ufold-8.6M* model with post-processing. The leftmost pair of violins show the F1 scores for the entire seen and unseen sets and the following violin pairs show the F1 scores for each constituent RNA family. The averaged F1 scores are shown at the top and also as dashed lines (white) in the corresponding violins. The values in the parentheses above the violins are the sequence counts in actual numbers (for the whole set or families with less than 1% population shares) or in percentages (for families with more than 1% population shares). Note that 23S rRNA only exists in the ArchiveII ND-L600 dataset and is thus shown as nan for the Stralign ND-L600 dataset.

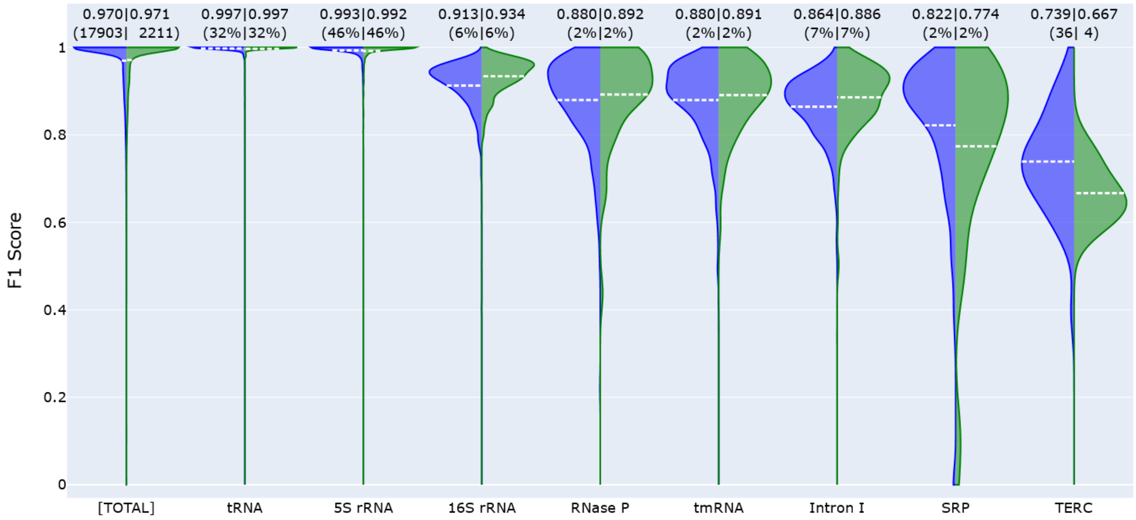


Figure S11. The F1 scores of the TR set (left, blue) and the VL set (right, green) for the SeqFold2D-1.4M model developed with the Stralign ND-L600 (Stral-ND) dataset randomly split into two subsets only: TR and VL. It is the same SeqFold2D-1.4M model shown in Fig. S9. The main observation is that no significant TR-VL variances (i.e., overfitting) exist on the whole or for individual RNA families. The order of RNA families shown follows the F1 scores of the TR set.

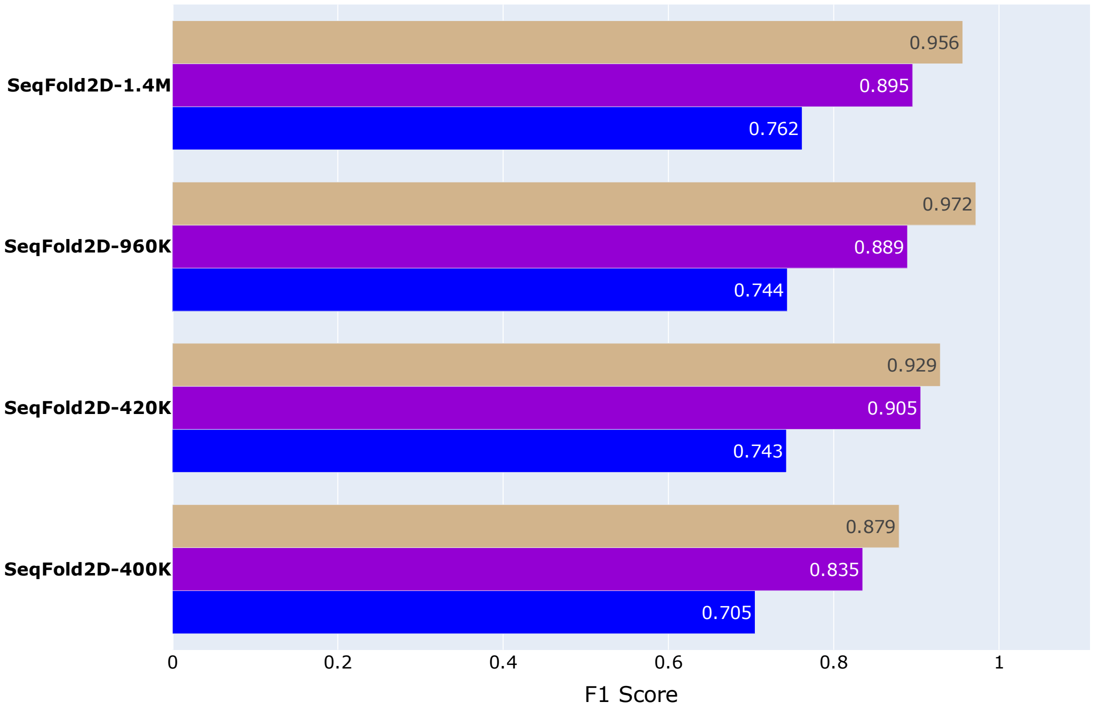


Figure S12. The F1 scores of the training (top, tan), validation (middle, violet), and test (bottom, blue) sets for the SeqFold2D models developed with the Stralign NR80-L600 (Stral-NR80) dataset randomly split into two subsets only: training (TR) and validation (VL). The test set is the Archi-Stral-NR80 dataset as presented in the main text. All SeqFold2D models for this cross-cluster study exhibit significant TR-VL variances (i.e., overfitting), while still attaining decent performances over the TS set. The two smallest models (400K and 420K) resulted from the explorations of depth vs. width and have design variables of (N=3, C=48) and (N=7, C=32), respectively. It is worth noting that increasing the number of parameters from 960K to 1.4M did not increase the performances on the TR and VL sets but resulted in slightly better performances on the TS set. We speculate that this may be due to the relatively small number of sequences in the TR set (2,653 total) that give a very rough gradient landscape for the algorithm to navigate, such that larger models can get trapped more easily in sub-optimal minima.

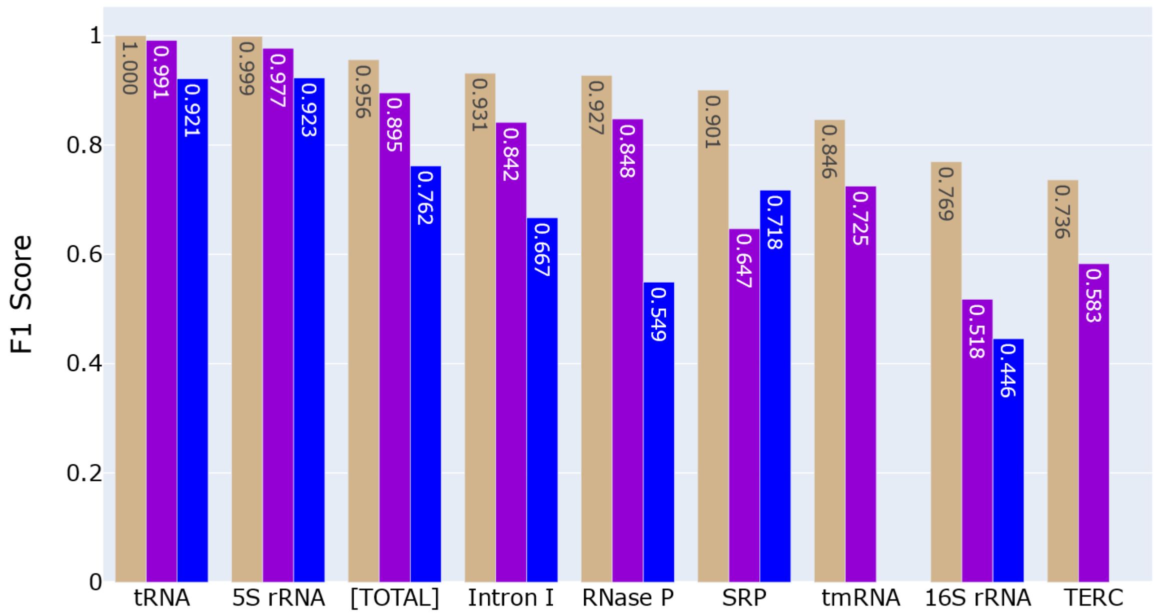


Figure S13. The F1 scores of the training (tan, left), validation (violet, middle), and test (blue, right) sets on the whole ([TOTAL]) and for individual RNA families for the SeqFold2D-1.4M model developed with the Stralign NR80-L600 (Stral-NR80) dataset randomly split into two subsets only: training (TR) and validation (VL). The TS set is the entire Archi-Stral-NR80 dataset. This is the same SeqFold2D-1.4M model shown in Fig. S12. The order along the x axis follows the F1 scores of the TR set. Note that the TS set does not have tmRNA or TERC sequences after removing sequences with above 80% similarity with the Stral-NR80 dataset. The main observation is that large TR-VL and TR-TS variances are observed for all RNA families and that the TR-TS variance is usually much larger than the corresponding TR-VL variance except for the SRP family.

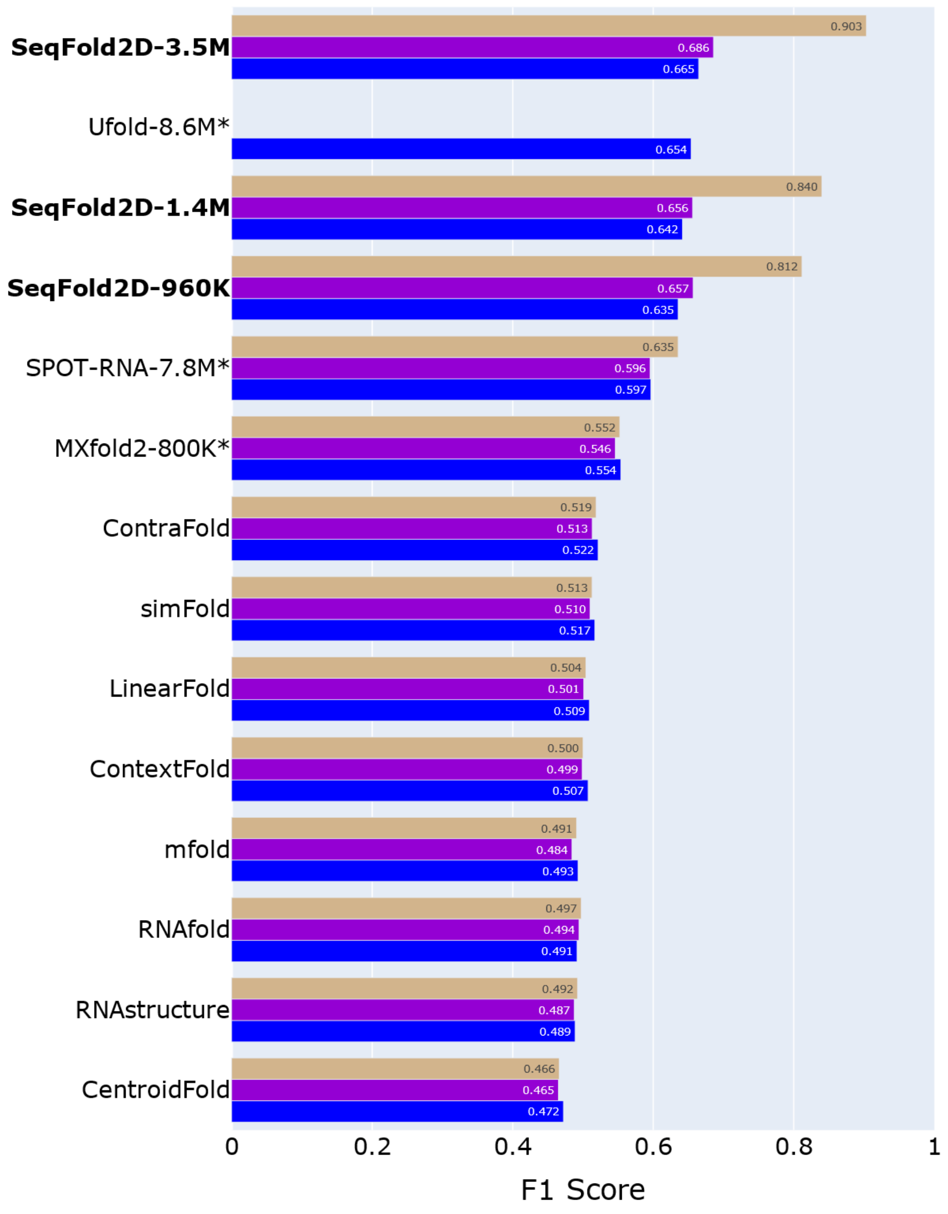


Figure S14. The F1 scores of the training (top, tan), validation (middle, violet), and test (bottom, blue) sets for three SeqFold2D models, a few other *de novo* DL models, and several traditional algorithms. Here the training, validation, and test sets are the bpRNA TR0, VL0, and TS0 datasets compiled by the SPOT-RNA team, respectively. The three datasets are expected to have independent, identical distributions, which are reflected by their comparable prediction performances by traditional algorithms. As discussed in the main text, the SeqFold2D models were trained to optimize the performance on the validation set, regardless of the magnitude of the train-validation variances. Ufold does not provide the saved model parameters trained with the bpRNA dataset, and thus only the value for the bpRNA TS0 set is available from the Ufold article (10). Note that the SeqFold2D models show even worse generalizability for the bpRNA-NEW dataset and we plan to use data augmentations techniques demonstrated by Ufold to improve generalizability in future work.

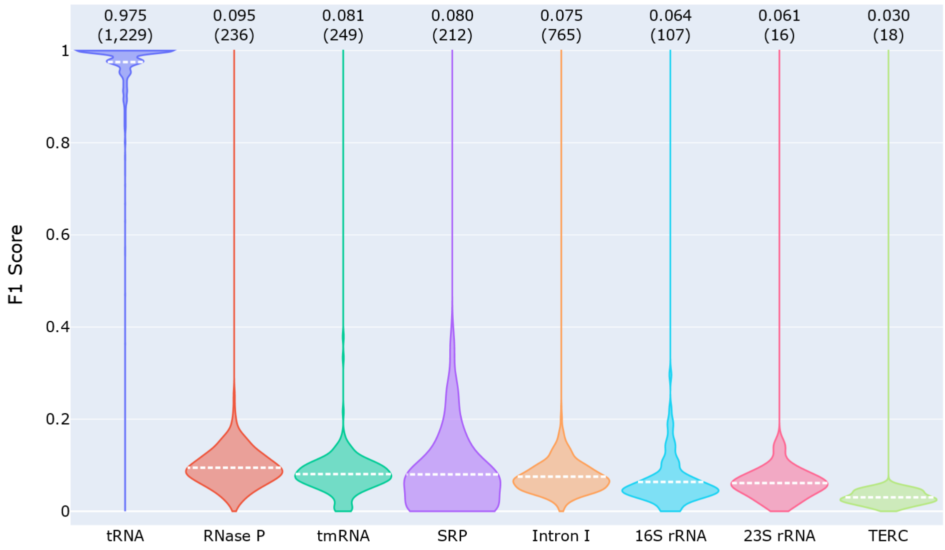


Figure S15. Cross-family generalizability for the SeqFold2D-400K model developed with the tRNA sequences in the Strive-NR80 dataset. Model training was stopped when the TR-VL variance became significant. While the model displays excellent performances over the seen sequences (the first violin), the performances over other family types fail completely.

### CONCLUSION AND DISCUSSIONS

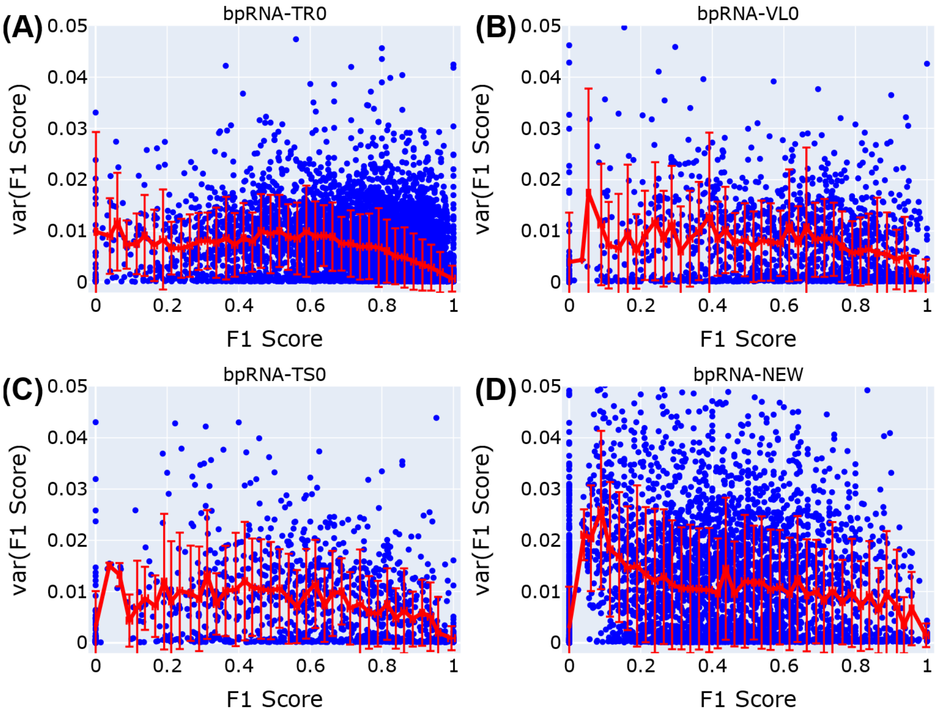


Figure S16. The correlations between the estimated variances and the actual values of the F1 scores on the training (A, bpRNA TS0), validation (B, bpRNA VL0), test (C, bpRNA TS0), and another independent test (D, bpRNA-New) datasets for the SeqFold2D-960K model.
